# Structured dynamics in the algorithmic agent

**DOI:** 10.1101/2023.12.12.571311

**Authors:** G. Ruffini, F. Castaldo, J. Vohryzek

## Abstract

In the Kolmogorov Theory of Consciousness, algorithmic agents utilize inferred compressive models to track coarse-grained data produced by simplified world models, capturing regularities that structure subjective experience and guide action planning. Here, we study the dynamical aspects of this framework by examining how the requirement of tracking natural data drives the structural and dynamical properties of the agent. We first formalize the notion of *generative model* using the language of symmetry from group theory, specifically employing Lie pseudogroups to describe the continuous transformations that characterize invariance in natural data. Then, adopting a generic neural network as a proxy for the agent dynamical system and drawing parallels to Noether’s theorem in physics, we demonstrate that data tracking forces the agent to mirror the symmetry properties of the generative world model. This dual constraint on the agent’s constitutive parameters and dynamical repertoire enforces a hierarchical organization consistent with the manifold hypothesis in the neural network. Our findings bridge perspectives from algorithmic information theory (Kolmogorov complexity, compressive modeling), symmetry (group theory), and dynamics (conservation laws, reduced manifolds), offering insights into the neural correlates of agenthood and structured experience in natural systems, as well as the design of artificial intelligence and computational models of the brain.

**Highlights:**

- Lie generative models are formalized using Lie pseudogroups, linking algorithmic simplicity, recursion, and compositionality with symmetry.
- Neural networks inherit structural constraints reflecting the symmetries in Lie-generated data.
- Similarly, agents, instantiated as neural networks tracking world Lie-generated data, reflect Lie structure and reduced-dimensional dynamical manifolds.
- Compositional structure in world data induces coarse-grained constraints, resulting in reduced manifolds that reflect the underlying generative process.
- Mutual Algorithmic Information (MAI) between the agent and the world emerges as shared symmetries in their dynamical interactions.
- These findings provide new insights for neuroscience, AI design, and computational brain modeling, emphasizing the interplay between data structure and agent dynamics.

## 1 Introduction

Understanding the fundamental principles of *agency* — spanning cognition, planning, valence, and behavior — and the shaping of subjective *experience* (perceived structure and emotion) remains a central challenge in neuroscience and artificial intelligence (AI), with broad socioethical ramifications beyond *homo sapiens*.^3,4^

In a series of papers^5–11^ we proposed a framework anchored in Algorithmic Information Theory (AIT) to study the phenomenon of *structured experience*. This term refers to the organized spatial, temporal, and conceptual structure of our subjective experience,^12^ encompassing both our perception of the world and our self-awareness as *agents* engaged with it, and serves as a critical bridge between first-person subjective experience and third-person scientific perspectives. Named after Kolmogorov complexity (𝒦), Kolmogorov Theory (KT) posits that a key function of the brain is information compression.^5,6^ This ability to compress information is equivalent to having access to a model of the world—unveiling its underlying structure — a critical foundation for cognition and the perception of structured reality.^7^

KT is related to and complements existing theories such as Active Inference and Predictive Coding and intersects with themes like the Information Bottleneck, grounded in Shannon Information theory.^9,13–15^ AIT is the generalization of Shannon information theory pioneered by Kolmogorov, Chaitin, and Solomonoff using computation theory (Turing).^11,16,17^ While Shannon’s entropy provides a probabilistic approximation to algorithmic complexity under noisy conditions,^11,17,18^ Kolmogorov complexity provides a universal, computation-theoretic lens whose role in defining and implementing model building has been highlighted in related domains, including AI and compression in Nature.^19–21^ These principles resonate with the longstanding pursuit of simplicity (low Kolmogorov Complexity), emphasized throughout history by thinkers such as Pythagoras, Plato, Aristotle, Epicurus, Occam, Leibniz (with special clarity in his *Discours de m*é*taphysique* (1696), as pointed out by Chaitin,^22^ Newton, Hume, Kant, and Einstein.^23^

*Algorithmic agents* are the central players in KT. They are defined as computational constructs using compressive world models, goals, and planning to interact effectively with their environment (Figure 2). Our definition of *agent* is inspired but not limited to natural agents. It characterizes them as information-processing systems equipped with features essential for evolutionary success: homeostasis (preservation of self) and, ultimately, *telehomeostasis* (preservation of kin). Examples of agents include animals, plants, and life in general, as well as properly designed AI systems combining a world modeling engine, an objective function, and a planning module.^8,9^

#### Definition 1.1.

An **algorithmic agent** is an information processing system with an objective function (goals) that interacts bidirectionally with the external world, inferring and running compressive models, planning, and acting to maximize its objective function.

Agent elements of especially relevance for our discussion are the *Modeling Engine* and the *Comparator*, which evaluates model prediction errors by comparing model-generated data with sensor data (see Figure 2). The existence of a *Modeling Engine* in the agent follows from the Regulator Theorem, a key result in the field of cybernetics that states that “Every good regulator of a system must be a model of that system”.^24,25^ For a regulator to effectively steer a system toward its goals, it must rely— implicitly or explicitly—on a model that captures the system’s relevant properties. In AIT terms, this requirement manifests as high mutual algorithmic information (MAI) between the agent and its environment, indicating the degree to which they share the underlying algorithmic structure.

KT is both a theory of cognition (behavior, third-person observations) and *experience* (phenomenology, first-person science), where the qualitative aspects of *structured experience* (𝒮) are associated with the algorithmic features of agent world models and their associated valence (𝒱).^10^ It hypothesizes that the *presence qualia* of structured experience in *algorithmic agents* emerge from the successful comparison of (compressive) model-generated data with world data^6,8,9^ and that the structure of all experience reflects the structure of models. Since successful models reflect structure in world data, KT predicts that the structure of experience ultimately stems from the structure inherent in the underlying *generative model* of world data. This raises the important challenge of characterizing the structure of world data and its corresponding agent models, with a focus on compositionally and recursion — key ingredients for compressive and efficient representations. But how is this expressed in dynamical terms?

The structure of models, central to KT, can be explored by instantiating the agent as a dynamical system. This approach is well-suited because computation can be understood as a dynamical process^27,28^—Turing machines can be interpreted as discrete dynamical systems characterized by transitions between states evolving over time. From this perspective, models and other components of the agent are programs that are “run,” and model and agent structure map directly onto dynamical structure,^9,10^ where dynamics refers to the time evolution of variables in the agent program. The connection between algorithmic and dynamical perspectives opens avenues for analytical methodologies. Tools such as bifurcation theory, differential geometry, and topology provide insights into algorithmic agents and the data they produce. These approaches are particularly relevant for exploring the brain state dynamics and attractor landscapes which can deepen our understanding of the mechanisms underlying neuropsychiatric phenomena and neurophe-nomenology,^29^ as depicted in Figure 1.

**Figure 1.**
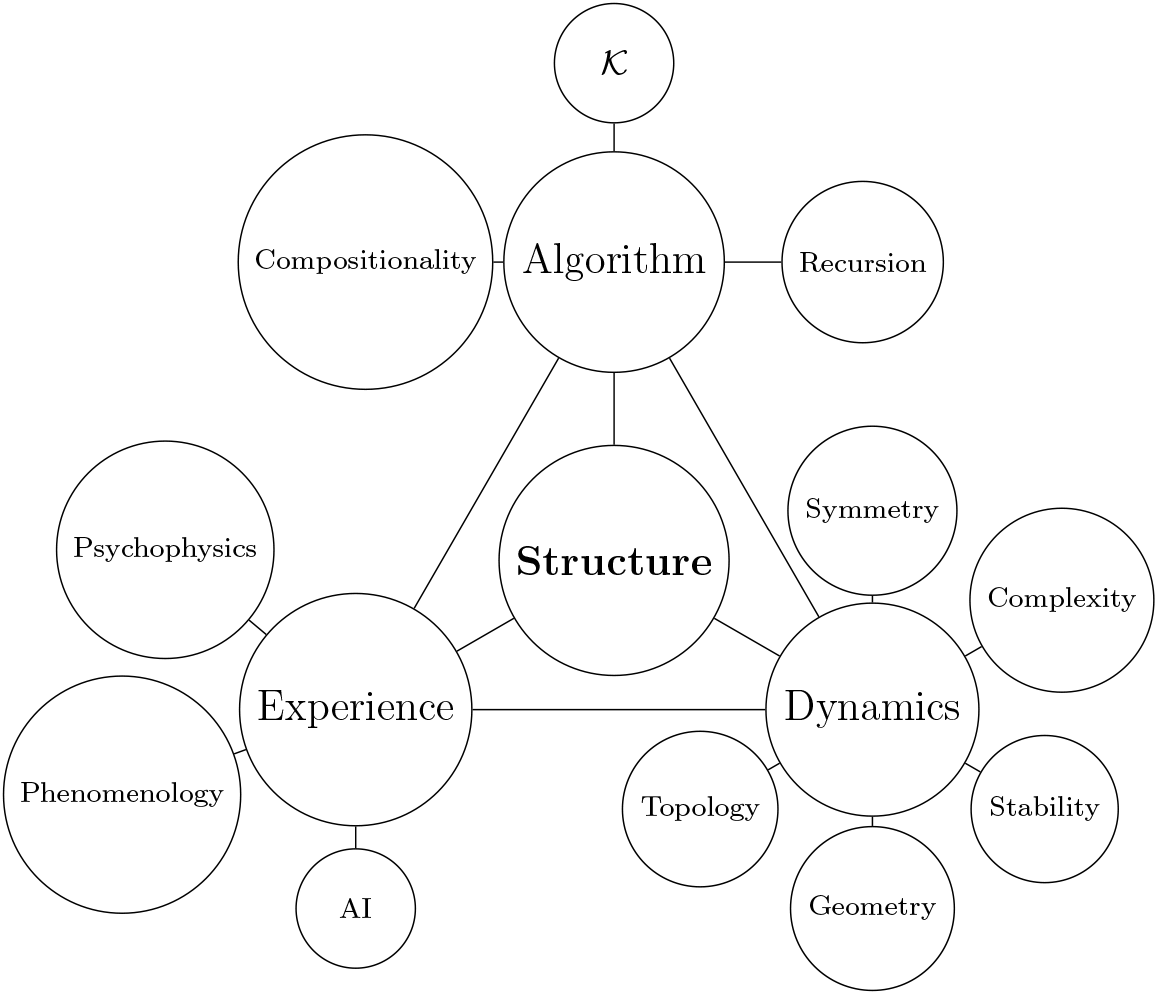
An illustration of the relationship between underlying mathematical model Structure, Algorithm features (such as algorithmic complexity 𝒦, i.e., program length, compositionally, recursion, cyclomatic complexity,^26^ etc.), Dynamics, first-person Experience (neurophenomenology), and in more detail some of the tools used in studying Dynamics, i.e., stability theory, bifurcation theory, chaos, geometry, topology, complexity (entropy, algorithmic), and symmetry, and Experience (including psychophysics, phenomenology and AI methods to characterize the structure of experience).

**Figure 2:**
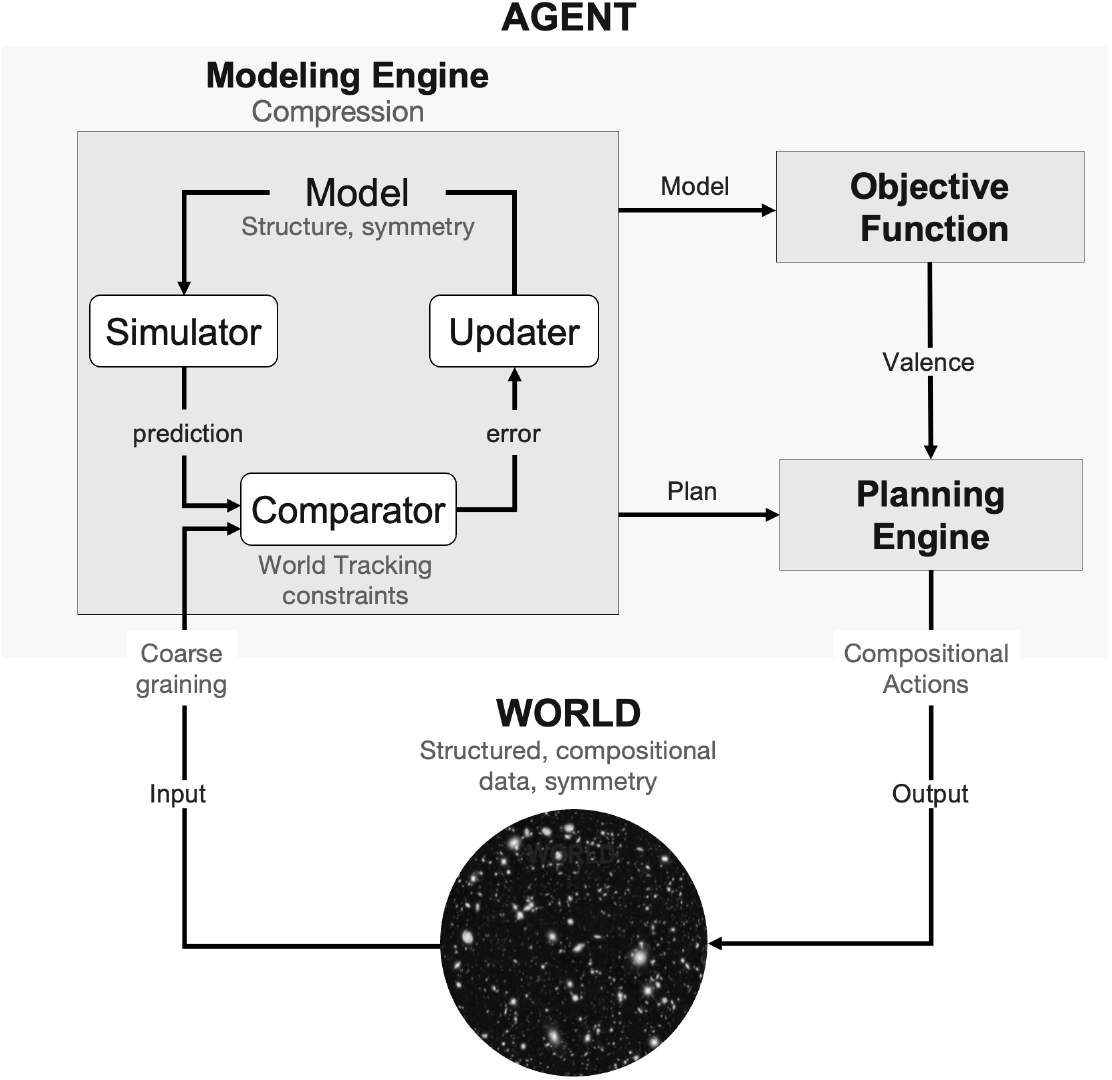
KT’s algorithmic agent, symmetry and dynamics. The **algorithmic agent**^8–10^ interacts dynamically with the World (structure, symmetry, compositional data). The *Modeling Engine* (compression) runs the current Model (which encodes found structure/sym-metry) and makes predictions of future (compositional, coarse-grained) data and then evaluates the prediction error in the *Comparator* (world-tracking constraint monitoring) to update the Model. The *Planning Engine* runs counterfactual simulations and selects plans for the next (compositional) actions (agent outputs). The Updater receives prediction errors from the Comparator as inputs to improve the Model. The *Comparator* is a key agent element monitoring the success of the modeling engine in matching input data. We reflect this process mathemat-ically as a world-tracking constraint on the dynamics (Equation 4.3, see also Section 4.4 and Appendix A.1).

Viewing the brain through the lenses of *computation* and *compression* provides valuable insights for both theoretical and empirical research. The computational viewpoint of brain function directly suggests that the brain operates as a dynamical system with unique computational and informational characteristics emerging at the critical points in complex systems.^28,30–33^ Criticality is increasingly recognized as a guiding principle for understanding the brain’s complex behavior^9,34–47^ (although some of its features, e.g., power-law neuronal avalanches, may stem from other dynamical mechanisms^48–50^). Neural systems are hypothesized to reach such states through evolutionary and developmental mechanisms driven by homeostatic plasticity.^29,51–54^

Regarding the role of compression, we have argued elsewhere that, from the perspective of natural agents, the world—defined as whatever is generating data on the other side of the agent—is, to a significant extent, simple and compressible.^8,9^ This assumption, which mirrors the “unreasonable effectiveness of mathematics”^55^ and Occam’s razor,^17,19^ underlies our rationale for the definition of agents as systems seeking to derive models of the world for survival, guided by a prior bias toward simplicity (compressibility, in the algorithmic information-theoretic sense).

AIT provides the foundational definition of compression: *Kolmogorov complexity* is defined as the length of the shortest program capable of generating a dataset (e.g., a stack of images of a hand^17^). As we discuss here, compression fundamentally relies on the presence of *symmetry* or *invariance*.^5^ The ability to compress a dataset is equivalent to having a model of the data, which implies the existence of patterns, invariances, or regularities that can be exploited.^7^ Intuitively, the length and structure of such a minimal program or model will be closely related to the symmetries within the dataset. A model that successfully compresses data essentially builds on and reflects the underlying symmetries or invariances. In this sense, the model *is* the invariant entity.

A variety of methodologies exist to characterize the structure of algorithms beyond traditional Kolmogorov-style measures. Classical software engineering metrics (e.g., cyclomatic complexity, Halstead metrics, and Henry–Kafura’s information flow) capture factors like control-flow branching, operator usage, and module coupling.^26,56^ In addition to these, one can examine *recursion depth* and nested compositionality,^57^ parallel or circuit depth,^58^ and category-theoretic notions of compositional hierarchy.^59^ Furthermore, functional and inductive approaches highlight structures such as iterated function composition^60^ and structural recursion,^61^ offering granular insights into how algorithms decompose data and transform it through layered abstractions.

While structure in algorithms manifests as *compositionality* and *recursion*, we expect that the dynamical systems arising from agent computations using these models—as observed in their structure, behavior, or neural activity—inherit and reflect the structure derived from the models. Consequently, by studying the relationship between the structure and symmetries of world data, the computational features of the models being run by the agent (model structure), the *structure* of neurophenomenological (first-person *reports*), behavioral, and physiological data (third-person records), can shed light on the hypothesis that an agent has (or at least reports) structured experience in a way that mirrors the structure of the compressive models it uses to track (structured) world data.

As part of this ambitious research program, we focus in this paper on *the relationship between the structure of input (World) data and the computational (constitutive, structural) and dynamical features of the successful world-tracking agent*. Through this analysis, we aim to provide a new perspective on phenomena such as changes in the complexity of neural data^29,42–44,62^ or the *manifold hypothesis*, which holds that natural data (including neuroimaging data) lies on lower-dimensional manifolds in its embedding space.^63–65^ We include insights from the compositional (hierarchical) nature of world data and neural processing and the associated necessity of coarse-graining.

Since invariance, structure, and symmetry are central to the problem at hand, we are naturally led to group theory and, more specifically, to the theory of continuous groups —Lie groups and their generalization. Next, we provide an informal definition of Lie groups and pseudogroups. Formal versions are provided in the Appendix A.2.

#### Definition 1.2.

A **Lie group** can be thought of as a set of symmetries that forms both a group (where we can combine symmetries and invert them) and a smooth manifold (so these operations behave nicely in a “smooth” sense). Concretely, its group multiplication (*g, h*) ⟼ *gh* and inverse *g* ⟼ *g*^−1^ are smooth maps.

In a **Lie pseudogroup**, we only require this structure to hold locally on a manifold. In other words, each transformation is only defined on a neighborhood within the manifold, but the usual group properties (identity, composition, inverse) still apply wherever these neighborhoods overlap.

The fundamental principles of symmetry and Lie groups, which have provided the foundation for modern physics, including relativity theory and quantum mechanics,^66,67^ are now being adopted in machine learning (see, e.g.,^68–72^) after pioneering work in the context of convolutional networks.^73,74^ This paper is a parallel development in the context of KT and, as we shall see, with applications in both computational neuroscience and machine learning.

In this paper, we provide a unifying framework to define and study *structure* across three domains: computation (via AIT), dynamics, and subjective experience. Specifically, we present a general model in which agents identify and exploit patterns in world data to build compressive representations, drawing on AIT as a foundation for compositionality and recursive structures.^17,18^ We link these AIT principles to invariance and symmetry, introducing a Lie Generative Model that connects these ideas with Lie group theory and pseudogroups.^1,75^ By showing how symmetries in the environment constrain the agent’s internal dynamics through a Comparator mechanism, we demonstrate that world-tracking conditions naturally induce reduced manifold structures in the agent’s state space.

The main novel contributions of this work are: (i) establishing a formal correspondence between compression-based theories of representation and Lie-theoretic approaches to invariance; (ii) presenting a unified Lie Generative Model framework that captures how environmental symmetries shape both the agent’s constitutive and dynamical structures; and (iii) illustrating how these links inform compositional and recursive perspectives on cognition and learning.

In the next sections, we start from the mathematical framework of Lie groups to formalize the concept of *structure* and provide a definition of *generative model*. This foundation allows us to explore how the algorithmic simplicity of a model is linked to representations of compositional or *hierarchical* Lie groups. We then demonstrate that neural networks, such as feedforward networks, inherently acquire structural constraints from the symmetry properties of the data they are trained on. Translating this to the context of recurrent neural networks (RNNs) leads us to discuss the central role of *symmetry* and *conservation laws*, which provide the basis for the concept of *compatible world-tracking constraints*. Building on this, we show that when a dynamical system aims to track the world, it is compelled to mirror the symmetries present in the data. This mirroring results in structural constraints and the emergence of *reduced manifolds*, which are lower-dimensional spaces that encapsulate the system’s essential features while respecting the imposed constraints. The compositional nature of world data gives rise to the notion of compositional or *hierarchical constraints and manifolds*. Our discussion is high-level and does not distinguish between natural and artificial neural networks, with implications for neuroscience, AI design, and brain modeling.

## 2 Generative models as Lie groups

Prior work in computer vision has highlighted the role of symmetries in image data,^76–78^ and in particular those associated with the group of rotations, translations, and dilations in *n*-dimensional Euclidean space. These can be considered as an extension of the Special Euclidean group *SE*(*n*) to include scaling transformations and is often referred to as the Similarity group, denoted as Sim(*n*).^79^ We recall that groups act on manifolds to effect transformations (see the Appendix for various definitions).

Artificial neural networks used for classification can be understood as representing invariance with respect to a *group of transformations*.^7^ This group defines a structure under which instances that are similarly classified form equivalence classes, capturing the shared features or invariances among those instances. This perspective offers a principled way to understand the representations learned by the neural network and the invariance properties it might acquire during training, potentially facilitating the analysis and visualization of the learned representations in terms of the group actions or guiding the design of network architectures. We aim to extend these ideas to more general *generative models*—the starting point in KT.

### 2.1 Classifying model-generated cat images

To make the discussion tangible but without loss of generality, we continue with the example of a classifier or autoencoder of cat images. We will work with several mathematical objects: the universe of all images ℝ^*X*^ (with *X* the number of pixels^a^), the subset of cat images (a very large or infinite set), a generative model of this set, and a symmetry group *G* transforming cat images into cat images.

#### Generative models

Our first assumption is that the space of cat images can be produced using a simple, smooth, generative model of cat images mapping points from a (relatively) low-dimensional manifold 𝒞 (the compressed, “cat image latent space”) to a larger image space—see Figure 3. This aligns with the view that the notion of “cat” is intrinsically a compressive model, an *invariance* encoded as a generative model using a few parameters. We formalize this in the following definition.

**Figure 3:**
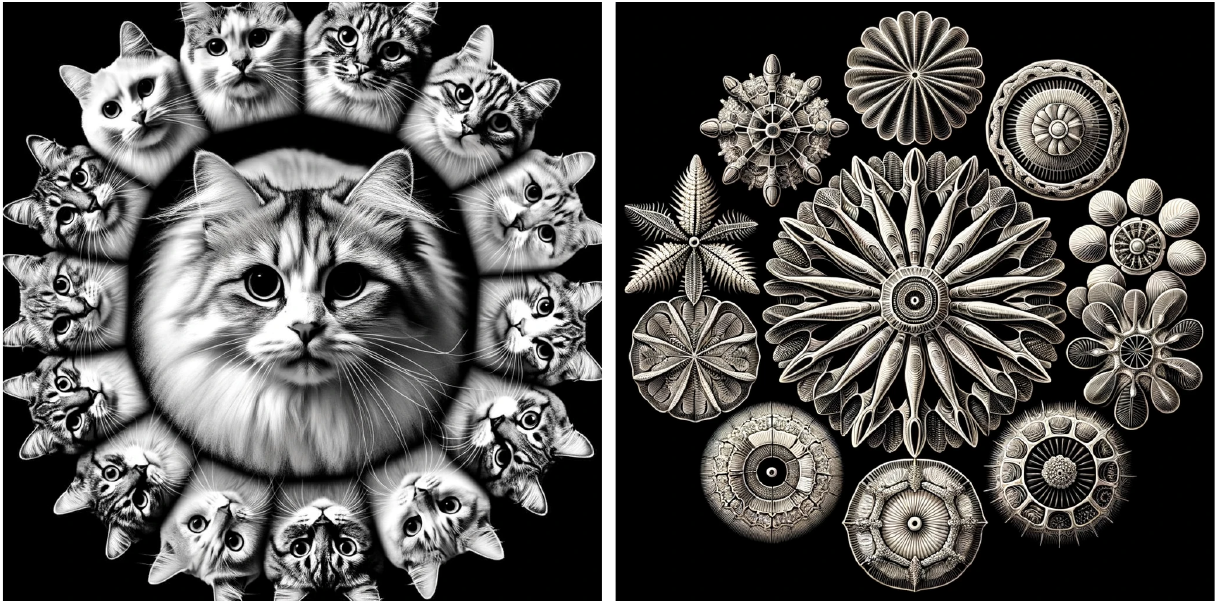
Variations from canonical cat (left) or diatom (right). An illustration of cat and diatom images (AI-generated in Ernst Haeckel’s style) derived from a central archetype (center) via the transformative action of a Lie group in latent space. In such a generative model, any image can be used as an archetype due to the transitivity of the acting group.

###### Definition 2.1.

By a *generative model* of images, we mean a smooth function mapping points in the M-dimensional configuration space manifold to X-dimensional image space, *f* : 𝒞 → ℝ^*X*^ so that *I*_*c*_ = *f* (*c*) ∈ ℝ^*X*^ for *c* ∈ 𝒞, with *M << X*.

We can locally represent the generative model using coordinates on the manifold, which we refer to as ‘patch’ coordinates, denoted by *θ* ∈ ℝ^*M*^, typically with *M << X*. In this context, *I*_*θ*_ = *f* (*θ*) ∈ ℝ^*X*^ represents the image generated by the model, where *M* is the number of parameters defining the model, and *X* is the number of pixels in the generated image *I*_*θ*_. Parameters are coordinates of cat space manifold 𝒞 (locally ℝ^*M*^) —cat image *configuration space*.

A subset of the space of all possible images corresponds to the category of “cats”. Suppose there is a group *G* that acts on the set of images of cats or, equivalently, their corresponding parameters in configuration (latent) space and transforms them into other images of cats, formally defined as the maps *G ×* 𝒞 → 𝒞, and *G ×* ℝ^*X*^ → ℝ^*X*^, corresponding to *γ*. *θ* = *θ*^*′*^ (configuration space) and *γ*. *I* → *I*^*′*^, with *γ* ∈ *G* (image space).

The largest group of transformations that can act on cat images while preserving their structure is the group of *automorphisms*, which in this case refers to all possible permutations of cat images.^7^ Within this large group, there exists a specific subgroup known as the diffeomorphism group, denoted as Diff(𝒞). This subgroup is an infinite-dimensional Lie group^80^ that consists of *smooth* (i.e., continuously differentiable) transformations or deformations of the configuration space manifold, mapping cat images to other cat images through the generative model.

To further formalize the notion of a generative model, we start from a reference image of a “cat” and define continuous transformations to other images of cats. A desirable property of a generative model defined this way is that it should not depend on the choice of a reference image—the concept of “cat” should not hinge on a specific canonical cat. To achieve this, we impose a *transitivity* requirement, meaning that the group of transformations must be able to link any cat image to any other cat image.

Since we can appeal to the infinite-dimensional group Diff(𝒞), we know we can always associate a Lie group with the generative model. However, to keep the generative model simple, particularly by ensuring it has a finite number of parameters, we make the stronger assumption: that the generative world model of cat images can, at least locally, be associated with a *finite* Lie group *G*.

Finite Lie groups capture the essence of continuous symmetries in a compact, finite-dimensional form, making them highly amenable for theoretical study and practical computation. In contrast to the general diffeomorphism group Diff(𝒞), which is often infinite-dimensional, finite Lie groups provide a structured approach to understanding global behavior from local information near the group identity element. This makes them particularly useful as candidates for simple generative models of world data, as they encapsulate key symmetries and invariances in a tractable manner through a few group generators. They also capture naturally the algorithmic notions of recursion and compositionality.

Since we can appeal to the infinite-dimensional group Diff(𝒞), we know we can always associate a Lie group with the generative model. However, to keep the generative model simple, particularly by ensuring it has a finite number of parameters, we make the stronger assumption: that the generative world model of cat images can, at least locally, be associated with a *finite* Lie group *G*.

Finite Lie groups capture the essence of continuous symmetries in a compact, finite-dimensional form, making them highly amenable for theoretical study and practical computation. In contrast to the general diffeomorphism group Diff(𝒞), which is often infinite-dimensional, finite Lie groups provide a structured approach to understanding global behavior from local information near the group identity element. This makes them particularly useful as candidates for simple generative models of world data, as they encapsulate key symmetries and invariances in a tractable manner through a few group generators. They also capture naturally the algorithmic notions of recursion and compositionality.

#### Recursion and compositionally in Lie group action

To illustrate this, recall that a small transformation generated by a Lie group element *γ* can be described as a perturbation around the identity element as *γ* = 1_*G*_ + *ϵT*, where *γ* is a group element representing a small transformation, 1_*G*_ is the identity element representing no transformation, *ϵ* is a small parameter representing the “size” of the transformation, and *T* is an element the Lie algebra, inducing infinitesimal transformations in the considered space (in our context, the “cat space”). This formulation offers a linear approximation of *γ* in the vicinity of the identity element, utilizing the Lie algebra structure derived from the Lie group.

Any element within the connected component of the identity can be represented as a product of infinitesimally perturbed identity elements, denoted by *γ* = 1+ ∑_*k*_ *ϵ*_*k*_*T*^*k*^,

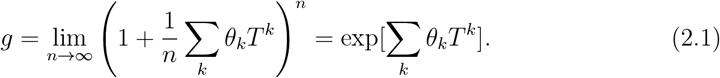

where *θ*_*k*_ are parameters and *T*^*k*^ are the generators of the Lie algebra.^75^ If *G* is a connected, compact matrix Lie group, the exponential map for *G* is surjective (covers the entire group).^75^ This *recursive* expression encapsulates the fundamental property of Lie groups that any group element can be decomposed into an (infinite) series of infinitesimal transformations. Compositionality extends this by including the sequential action of different group elements. In algorithmic terms, recursion is a for loop, and composition is a set of nested for loops.

#### Lie pseudogroups and generative models

However, in a topologically complex latent space, finite Lie groups (globally defined transformations) may be too restrictive as they require a single structure acting globally. A Lie *pseudogroup*, by contrast, focuses on partial transformations with restricted domains that can be iteratively composed when their domains overlap, providing both local flexibility and global reach. This is particularly suited for “walking” the latent space of a generative model in iterative steps—chaining local diffeomorphisms across patches in a manner that respects the underlying manifold structure. Consequently, a pseudogroup formalism often emerges as the most robust solution to handling high-dimensional, topologically intricate configuration spaces

This motivates the following definition, closely related to the notion of a group of transformations acting on a manifold (see the Appendix):

###### Definition 2.2.

An *r*-parameter generative model *I* = *f* (*c*), *c* ∈ 𝒞, is a *(Lie) generative model* if it can be written in the form

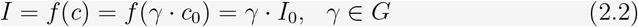

where c_0_ is an arbitrary reference point, *I*_0_ ∈ ℝ^*X*^ is an arbitrary reference image, *f* is a smooth function, and *G* is an *r*-dimensional Lie pseudogroup.

For example, in the case of cat images in a particular manifold patch, we may write *f* (*θ*) = *h*(*γ*_*θ*_ · *P*_0_) = *γ*_*θ*_ · *I*_0_, where *h* is a projection function (describing the operation of a camera, which, for simplicity, we may assume to be fixed, with uniform background, etc.), and *P*_0_ is an arbitrary reference point-cloud capturing the 3D form of a particular reference cat. In the second equality, *I*_0_ is the corresponding reference cat image, and *γ*_*θ*_ is a representation of the group *G* on that object. The action of the group element *γ* varies depending on the type of object it is applied to—whether a point cloud, an image, or a parameter—resulting in different *representations* of the group for the same transformation. For instance, when dealing with point cloud data and rotations, the group’s action can be described by matrices, giving us a linear representation in a vector space. However, linearity of representations may not generally hold—the representations found in natural settings are often non-linear, outside the domain of classical *representation theory*.^1^ Thus, although we refer to the map from the latent (or object) space to sensor measurements (e.g., images) as an “induced representation,” it may generally be a non-linear group action—and, due to the non-injectivity of the sensor projection, it may also be multi-valued.

The function *f* must be smooth to leverage the recursive and compositional nature of the Lie action, ensuring that the resulting generative model is compressive (e.g., it cannot be a giant look-up table). This connection ties the structure of the Lie generative model to the notion of simplicity as defined by Kolmogorov complexity in algorithmic information theory.

A Lie pseudogroup structure enables the navigation of the configuration space manifold in a succinct manner. By using Lie groups, models can capture the essential characteristics of objects in a mathematically rigorous and computationally efficient manner. This approach facilitates both the understanding of the learned representations in neural networks and the practical implementation of generative models.

The assumption of finite dimensionality is not restrictive when the generative model depends on only one continuous parameter. This applies to processes associated with rotations, translations, and dilations since these correspond to finite Lie groups. It also holds for transformations of *shape space*, e.g., if the cat image generator source is a robot with any number of rigid joints, which can be described using the special Euclidean group SE(*n*).^81^ More broadly, while it is not always clear whether images generated by models like generative neural networks or 3D modeling programs (e.g., Blender^82^) possess the group structure of a finite Lie group, generative models exhibiting transitivity, smoothness, and a finite number of parameters may often be described by a finite Lie pseudogroup.^78,83,84^ We discuss the limitations of the finite-dimensional assumption in Appendix A.5.

In summary, data produced by a generative model can always be locally generated by a finite-dimensional Lie group. Some generative models admit global finite-dimensional Lie pseudogroup actions and are, therefore, Lie generative models in our definition. This will hinge on the geometry of the generative model configuration space 𝒞. It is nevertheless always possible to locally cast generative models as Lie generative models.

### 2.2 Implications of invariance under group action

To illustrate a Lie recursive and compositional generative model and study the implications of invariance, imagine generating cat images by taking pictures of a jointed cat robot (see Figure 4) for training a deep neural network classifier. In this context, *invariance* means that when we apply transformations, such as rotations or pose changes that do not alter the semantic identity of the object, the output of the model should remain unchanged—i.e., the classifier should still recognize the object as “cat.” The robot’s varying state is expressed through generative compositionality and is described by the Product of Exponentials formula from robot kinematics,^85^

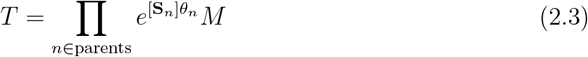

where *T* ∈ *SE*(3) is the final position and orientation of the end effector (e.g., a particular cat claw) in the special Euclidean group, *θ*_*n*_ is the vector of joint variables for the *n*-th joint, [**S**_*n*_] is the skew-symmetric matrix representing the screw axis of the *n*-th joint, 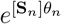 is the matrix exponential representing the action of the *n*-th joint’s transformation, and *M* is the initial (home) configuration of the end effector.

**Figure 4:**
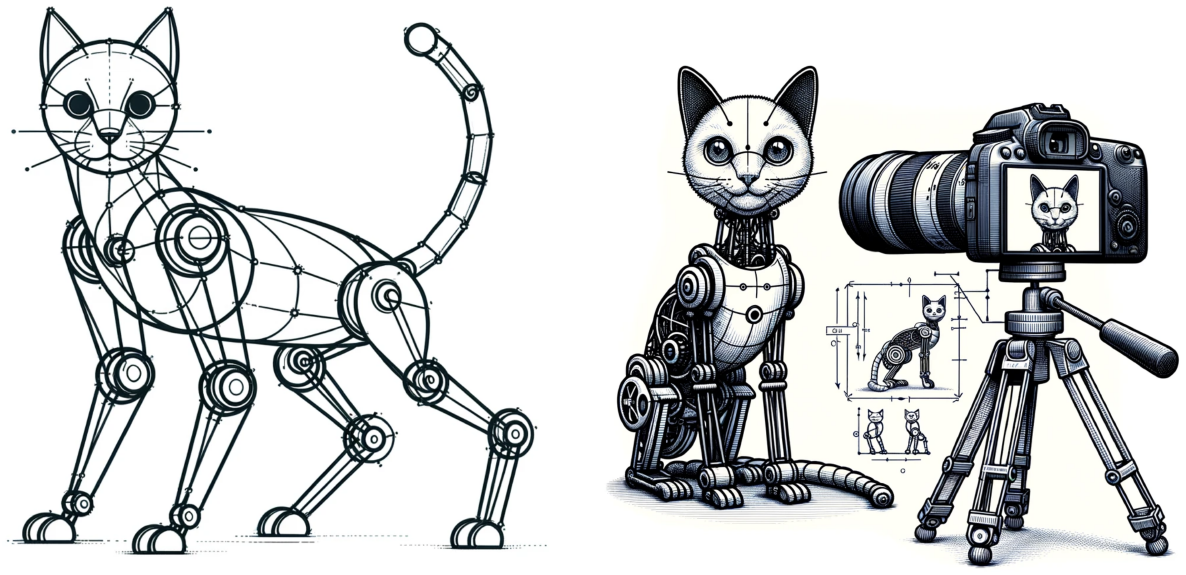
Cat robot and generated cat image. A representation of the generative model of a cat derived from a robotic construction using the product of exponentials formula (Eq. 2.3) in robot kinematics. The robot consists of a set of joints controlled by a Lie group. Left: cat robot using joints to control pose and expression. Right, projection into an image using a camera. (These images are themselves AI-generated).

**Figure 5:**
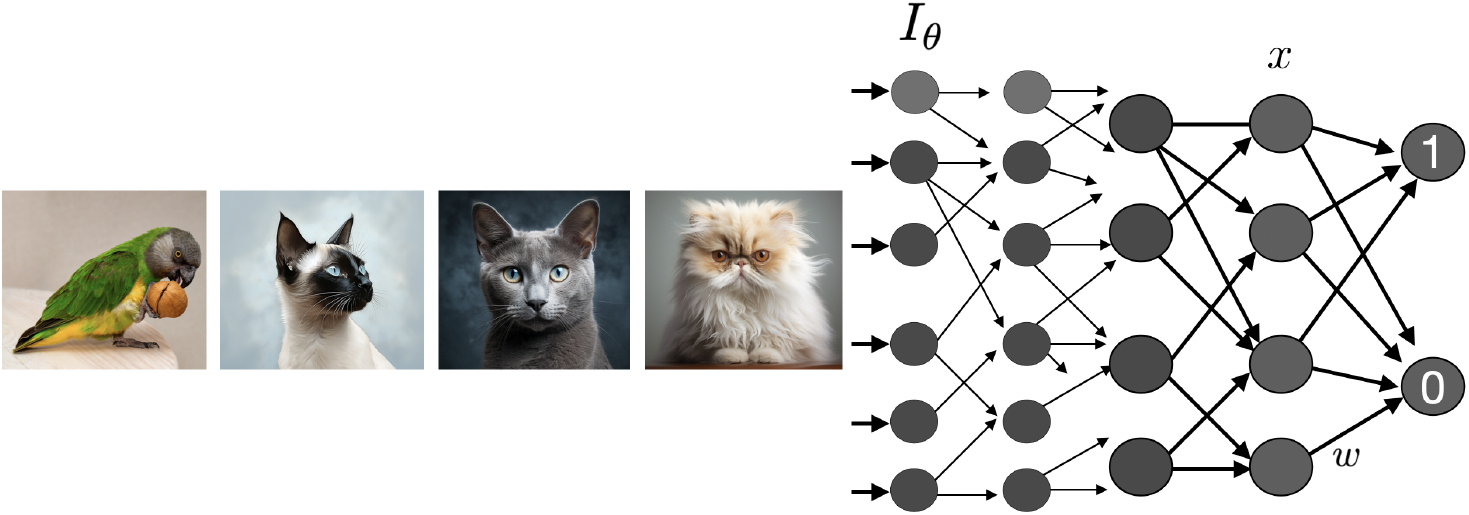
Image classification task. Classification of cat images can be seen as learning the invariances of a generative model. Such a network can also be implemented using an autoencoder of cat images with a skip connection to detect anomalies.

**Figure 6:**
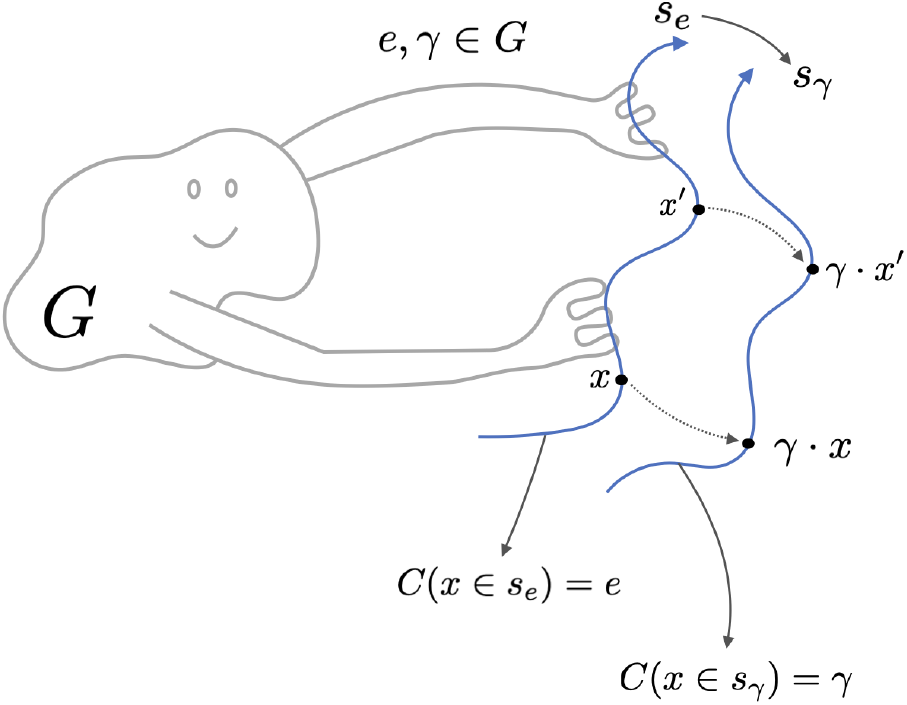
Group acting on solutions space and the (generalized) Noether’s theorem. The action of a group on ODE trajectories can be used to provide a labeling system for them. A reference solution trajectory *s*_*e*_(*t*) (*e* is the identity element in G) is moved to *s*_*γ*_ under the action of element *γ* ∈ *G*. This gives rise to conserved quantity *C*(*s*), which is labeled by group elements (functions from phase space to group elements after the choice of some reference solutions—convention). [Figure inspired by the illustration of the action of the Galois group on polynomial equation solutions in *Tom Leinster’s Galois Theory (Fig 2*.*1)*.^96^]

Here, we assume that SE(3) is realized as a matrix Lie group, so the Lie algebra se(3) becomes a finite-dimensional vector space of 4 × 4 matrices (with a specific block structure). Accordingly, each joint’s screw axis [**S**_*n*_] lies in se(3), and the map 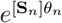 is the usual matrix exponential from the Lie algebra to the group. In this sense, we move from the manifold description of SE(3) to its vector-space representation (the Lie algebra) for the sake of computational convenience.

In this sequential composition, the order of joint transformations is crucial and reflects the non-commutative nature of the Lie group *SE*(3). Each joint’s transformation is applied in a fixed sequence, ensuring that the resulting configuration accurately represents the compounded effect of all joint movements. This ordered product does not assume that the individual transformations commute; instead, it leverages the inherent structure of the Lie group to maintain the correct spatial relationships between joints.

To represent the full state of the robot, a similar expression is applied to each joint and, by extension, to the entire robot. Each joint corresponds to a representation of a Lie group, and the calculations—including those involving the generators—are compositional. Leveraging this Lie-group compositional structure, we can generate an arbitrarily large yet highly compressible dataset of robot pose images.^7^

This principle may extend to other generative models of the world, such as more realistic images of cats. Fundamental physical laws often exhibit symmetry principles, such as covariance in general relativity or gauge invariance in quantum field theory. While it is not immediately clear how these symmetries propagate after the coarse-graining of phase space by an agent’s sensory system, it is plausible that they persist in some form. Alternatively, natural selection may favor agents that coarse-grain the world in a way that allows them to exploit compositional representations.^11^

Our network classifier aims to output “1” if an image contains a cat and “0” otherwise. While the universal approximation theorem^86^ guarantees that a sufficiently large network can approximate this classification function, it does not address how efficiently such a network can learn. Neural networks achieve this task efficiently, as shown in,^87,88^ if (1) the data is *compositional*, and (2) the architecture leverages that compositionality through a sufficiently *deep* structure. The former hinges on the presence of a succinct generative model—akin to having a “short program” (recursive, compositional) in algorithmic information theory. The latter leads to the design of deep or truly recursive architectures. The same reasoning applies to compressive autoencoders, where exploiting compositionality enables learning compact latent representations.

The equations for a general feedforward neural network are

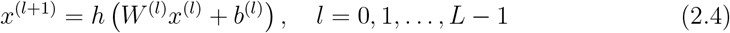

where *x*^(*l*)^ is the vector of neuron activations at layer *l, W* ^(*l*)^ and *b*^(*l*)^ is the weight matrix and bias vector at layer *l*, respectively, *h*() is the activation function applied element-wise, and the input data is fed as *x*^(0)^. The equations can be compactly expressed as *x* = *h*(*Wx* + *b*), where *x* is a vector encompassing the activations of all nodes in the network, structured in a way that respects the layered organization of the network and *W* is a block matrix describing the connections between the nodes, including the weights associated with each connection.

We require the classifier to be invariant under the action of the group *G*, whether it is finite or an infinite Lie group. Specifically, if the network classifies a point *θ* ∈ 𝒞 (via the image *I*_*θ*_ generated using parameters *θ*) as a “cat,” it should also classify the point *γ* · *θ* as a “cat” for any *γ* ∈ *G*. If the group is a finite Lie group, it is sufficient to study the *finite* number of generators of the group—linear operators in the group’s Lie algebra. An important property of Lie groups is that any element can be expressed in terms of actions near the identity element, although this applies only to the part of the group connected with the identity. The composition of group actions from elements near the identity covers the connected part. The recursive nature of Lie groups ensures that to establish invariance or equivariance of a function under group action, it is sufficient to verify this property for the generators *T*_*k*_. The behavior under infinitesimal transformations near the identity informs the behavior under the full group action. However, for disconnected Lie groups, invariance must be confirmed for each component separately.

A change in the input image from a class-preserving infinitesimal transformation will propagate through the network and affect the state *x*. The transformation can be described by *ϵ* where *x* → *γ* · *x* = *x* + *δx*, with *δx* = *ϵ Tx*, where *δx* is a small change in the state vector *x, T* is a linear transformation generator associated with the group action, and *ϵ* is a small parameter. This induces the transformation

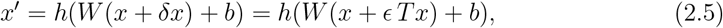

and we seek conditions under which the output of the network is invariant to first order in *ϵ*.

Using the chain rule, we can express the transformation matrix at a given layer *l* in terms of the transformation matrix at the input layer and the Jacobian matrices of the transformations at all preceding layers,

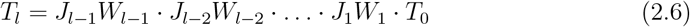

where *J*_*i*_ denotes the Jacobian matrix of the transformations at layer *i*, encompassing the activation functions, and *T*_0_ is the transformation matrix at the input layer, the linear approximation to *I*_*θ*_ = *γ*_*θ*_ · *I*_0_ ≈ *h*((1 + *ϵ T*_0_)*θ*_0_).

A necessary (but not sufficient) condition for the invariance of the neural network output with respect to the group action is that there exists at least one layer *l* for which the condition *J*_*l*_ · *W*_*l*_ · *T*_*l*_*x*_*l*_ = 0 holds for all cat image inputs (see^78^ for similar reasoning on the conditions imposed by symmetry). This ensures that the transformed state vector *x*_*l*_ + *T*_*l*_*x*_*l*_ remains in the null space of the weight matrix *W*_*l*_ at layer *l*, guaranteeing the invariance of the state of this layer with respect to the transformations generated by the group action. In other words, the space of cat images must be in the kernel of the operator *T*_*l*_ for some layer *l*. This property propagates to higher layers, ensuring the network’s output remains invariant to these transformations.

For the feedforward autoencoder, the analogous *equivariance* requirement is that the transformation of the output matches that of the input, namely,

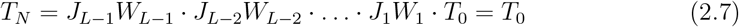

when acting on cat images. This sets an *M* -dimensional family of constraints on *W* requiring that the matrix *T*_*N*_ = *J*_*L*−1_*W*_*L*−1_ · *J*_*L*−2_*W*_*L*−2_ · … · *J*_1_*W*_1_ has the space of cat images as an eigenspace corresponding to the unit eigenvalue. These are the necessary requirements for invariance or equivariance under transformations.

Systems specifically designed for invariant behavior, such as convolutional networks with pooling layers, explicitly implement some of the required symmetries. Generalizing this idea involves designing networks that “filter” data across group orbits (e.g., such as translations or rotations), then analyze the results (e.g., computing the maximum or the average),^69^ or using data augmentation techniques, or finding fundamental architectural principles for group invariance.^89^ The success of hierarchical, recursive, deep architectures reflects this principle.^63,68^

In summary, imposing invariance of the classification of data produced by a generative model leads to equations that constrain the weight parameter matrix under transformations associated with the generative model. This approach is valuable for designing neural networks—where the weight equations can be enforced in the loss function during training—provided the generators are known.

Conversely, these principles can be useful for inferring symmetries underlying world data from the structure of networks trained on it.

## 3 The agent as a dynamical system

Although feedforward networks can be viewed as dynamical systems, their computational capacity is constrained by a fixed number of sequential evaluation steps. A broader perspective, bridging both algorithmic and biological paradigms, arises from the use of ordinary differential equations (ODEs), which have the potential to instantiate universal Turing machines.^90–92^ Differential equations can be interpreted as recursive neural networks (RNNs), modeling continuous dynamics similarly to recursive processes in NNs. This analogy becomes particularly clear when the ODE equations are discretized.

While the algorithmic perspective defines the agent as a program or Turing machine exchanging information with the environment, the dynamical view describes it less abstractly (but equivalently) and in a manner that makes more clear its connections to physics and biology—e.g., as a human brain or a neural network running on hardware. As a dynamical system, the agent is characterized by state variables and rules governing state transitions as a function of an external parameter denominated as *physical time*.

We define some terminology used in what follows. The *phase space* in dynamical systems theory is a multidimensional manifold (see Appendix for mathematical details) where each point specifies the state of the system (the necessary and sufficient information to project the state in time). Finally, by the *geometry of the phase space*, we refer loosely to the structure and properties of the subspace of dynamical trajectories in space, including the presence of any special points or areas, such as *attractors* or *repellors*.

### 3.1 General model

A general model of the state of neurons or other elements of a dynamical system receiving external inputs can be written in the form of a multidimensional ordinary differential equation (ODE),

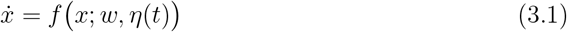

with *x* ∈ ℝ^*X*^ is a highly dimensional state variable representing the activity of neurons or neuron populations (e.g., membrane potential or firing rate) and where *w* stands for connectivity parameters (and maybe others). For example, in a model of the human brain, each coordinate may represent the average firing rate of populations of neurons (a neural mass model) or a more detailed ternary code of silences, spikes, and bursts in each cell.^93^ This finite-dimensional equation can be generalized to a partial differential equation (a neural field equation), but these details are not essential in what follows.

The input term *η*(*t*) reflects inputs from the world represented as an external forcing independent of the agent’s state that makes the equations non-autonomous (i.e., explicitly time-dependent). We can imagine it representing random kicks to the trajectory or a more steady, purposeful forcing received by agent sensors (we provide a generalization of this model to include inputs and outputs reflecting interaction with World dynamics in Appendix A.1). External inputs—which, from the perspective of a Universal Turing machine, can be viewed as reprogramming the agent—can force changes in the state trajectory or even drive the system’s landscape through dramatic topological transitions.^29,94^ For instance, such changes in topology can be induced by time-varying parameters associated with bifurcations in the system.

We call Equation 3.1 the “fast time” dynamics equation,^29^ as it governs dynamics at short time scales (seconds), where connectivity parameters *w* are assumed to be constant (in reality, *w* may be dynamic at slower time scales, reflecting *plasticity* phenomena). This equation fixes a *dynamical landscape*, the geometry of trajectories in phase space defined by the set of ODEs. More precisely, the landscape is the manifold defined by the motion of trajectories in the phase space manifold (with local coordinates *x* ∈ ℝ^*n*^). The dynamical landscape is determined by the form of the function *f* () and parameters *w*.

While we focus on deterministic ODEs for clarity in this work, future research will explore the implications of explicitly modeling *η*(*t*) as a stochastic process. The external input *η*(*t*) in our model can be extended to stochastic processes, leading to a stochastic differential equation (SDE) framework. For instance, *η*(*t*) could follow an Ornstein-Uhlenbeck process, introducing biologically realistic noise into the dynamics. This extension allows the exploration of noise-induced phenomena, such as stochastic bifurcations, attractor transitions, and resonance, adding richness to the system’s behavior.

### 3.2 Conserved quantities and symmetries

Let us focus now on the case where *η*(*t*) is constant or very slowly varying compared to the other dynamics. In the context of dynamical systems, *symmetry* is a transformation that maps a solution of the ODEs into another or, equivalently, maps the set of solutions into itself. More precisely, by the symmetry of an ODE, we mean that there is a group *G* such that for all elements *γ* ∈ *G*, if *x* is a solution to the equations, then *γ* · *x*, a transformation of *x* specified by the group element, is also a solution to the ODE.

Since the only indeterminacy in the solutions to the ODEs are the initial conditions, we can also view symmetries as the action of a group on the initial conditions manifold (integration constants). Symmetries partition the solution space (or initial condition manifold) into orbits under the group action. If the group acts transitively on that space, then, indeed, one can map any solution to any other. Otherwise, solutions lie in distinct orbits, and only solutions within the same orbit can be mapped into each other. Identifying symmetries in a system simplifies its analysis by organizing solutions into equivalence classes under the symmetry group. Because solutions within each class can be mapped into one another by the group action, it suffices to study one representative solution per class, with the group element labeling how solutions transform within that class.

Symmetries can be categorized as discrete or continuous, and in the latter case they are described by local Lie groups. Paralleling the implications of Noether’s theorem,^66,67,95^ the existence of symmetries can help in analyzing and solving ODEs. For example, the existence of a continuous symmetry means that we can locally define a coordinate system in which some of the coordinates do not appear in the equations other than in the derivatives (they are cyclic).^95^ Furthermore, there is a deep connection between symmetries and conservation quantities, as we now describe.

A *conservation law* is a map from solutions to a manifold. For example, energy conservation is a map from solutions to Newton’s equations to real numbers. Momentum conservation is a map from solutions to ℝ^3^.

The relationship between symmetry and conservation laws can be understood as follows. If a system is invariant under the action of a group *G*, then for any solution *x*(*t*), applying a group element *γ* ∈ *G* yields another solution, *γ* · *x*(*t*). Consequently, one can label every solution by the group element needed to map a reference solution (corresponding to the identity element *e* ∈ *G*) to it.

Under standard uniqueness conditions (guaranteed by the Picard–Lindelöf/Cauchy–Lipschitz Theorem^97^), trajectories in phase space do not intersect—apart from the trivial case of time-translation symmetry in autonomous ODEs. Hence, each point on a trajectory can be uniquely associated with a specific group element (disregarding temporal translations in autonomous systems). This induces a mapping from phase space to the group that remains constant along each trajectory. That constant constitutes the conserved quantity associated with the system’s symmetry. This construction parallels Noether’s statement: *continuous symmetries imply conserved quantities*.

Thus, we can think of this process as the act of creating a *structured labeling system* for the solution space and then phase space, using the group. *The group labels, as the output of functions from phase space to group elements, are the conserved quantities*.

To illustrate this, consider the equation 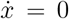 with *x* ∈ ℝ. It displays translational symmetry: if *x*(*t*) is a solution, then 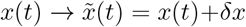 is also a solution. The group’s action is to translate solutions along the *x*-axis (phase space is 1D in this example). The conserved quantity is *C*(*x*) = *x* = *x*_0_, which is shifted by the action of the group, *C* → *C* + *δx*. We provide more simple examples of symmetry transformations that shift associated conserved quantities in Appendix A.6.

Finally, a well-known example in classical mechanics is the Kepler problem (two-body problem in celestial mechanics), an autonomous ODE in six-dimensional phase space, where a (bound) solution may be classified using conserved quantities corresponding to symmetries: the shape of an ellipse (eccentricity), its size (the energy or total angular momentum, one parameter), and the orientation of the plane containing it (three parameters). The latter results from the rotational symmetry of the problem and allows for the classification of the solutions with the orientation of the plane (the angular momentum). We can think of the specification of a solution in a constructive way: specify first the energy (one parameter) and eccentricity (one parameter) of an ellipse with its major axis along the *x*-axis and a minor axis in the *y*-axis. Then, rotate this plane in 3D (three parameters from the SO(3) or, equivalently, the SU(2) group). The existence of these five conserved parameters in the Kepler problem or in other similar central potential problems stems from symmetries of the equations (the O(4) group).^98^

More generally, *N* -dimensional ODEs have *N* − 1 conserved quantities (essentially functions of the constants of the motion, see Appendix A.6). An ODE with constraints (an algebraic differential equation) has solutions if the constraints are functions of the constants of the motion (and hence constant). The conserved quantities can be used to create, at least locally, new coordinates, which are then constant and act as (group) labels of solutions. The symmetry of the equation is associated with transformations that shift these constants.

## 4 The world-tracking condition

Because of the time-dependent input term, it appears difficult to say very much about the behavior of the fast-time equation (3.1). However, there are two important elements of the model that we can leverage. Namely, we assume that:

1. Agent inputs are generated by simple rules (the world is inherently simple), i.e., by a hierarchical generative model as discussed above.
2. The agent is able to “track” its inputs, i.e., uses an internal generative model to approximately match them.

We now analyze the mathematical consequences of these statements.

First, the assumption that world data is the product of a simple generative model—as discussed in the previous sections—implies that the constrained dynamics of the agent can be studied from the point of view of symmetry. We may think of the external world as providing stochastic but structured input to the dynamic system.

Second, one of the objectives of the agent is to track world data through the implementation of the world model in its modeling engine, which is to be compared with the data (this task is carried out by the *Comparator* in the KT agent model, see Figure 2).

To make this comparison meaningful, there must be a readout from the agent’s internal state that matches the tracked world data. This necessity imposes a dynamical constraint on the system: the agent’s internal states must evolve so that their readouts correspond to the external inputs. This constraint ensures that the agent’s internal dynamics are aligned with the environment, enabling accurate tracking and adaptation.

As we will see, the implications of world-tracking are, in fact, twofold. On the one hand, the dynamics of the system are constrained to lie in reduced manifolds. On the other, the architecture of the equations implementing world tracking needs to satisfy some requirements, as we already saw above in the simpler case of a feedforward classifier.

### 4.1 Constrained dynamics I

We begin with a simple toy model to study the role of symmetry and conservation laws. Consider the constrained system of differential equations (or *differential algebraic equation*^99^),

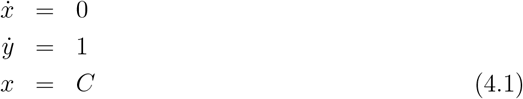

The first two equations are regular ODEs, and the last is a constraint (*C* is an arbitrary parameter), a toy world-tracking condition. These equations happen to be solvable for any value of *C*, because the general solution to the ODE subsystem is

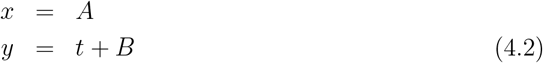

Thus, it is possible to select the *integration constants A* and *B* that satisfy the constraint. This occurs because the constraint is not independent of the conserved quantities of the system. The same conclusion follows if the third equation in 4.1 is replaced by *f* (*x*) = *C*, for any function *f* such that *C* is in its range.

Another way to view the compatibility of the equations is that the ODE subsystem displays translational symmetry in *x*, with the symmetry *x* → *x* + *ϵ*. This means that if *x* is a solution to the differential equations, then so is *x* + *c*, for any constant *c*. Hence, we can shift the trajectory *x* by any value we wish and still obtain a valid solution to the ODE.

Conserved quantities in and ODE, such as *x* = *C* above, can be used to create new coordinates with zero derivatives, which manifest the symmetry, just like *x* in our example. The symmetry group shifts solutions across the conservation surface, i.e., it shifts the constant associated with the conserved law. Further examples are provided in Appendix A.6.

### 4.2 Constrained dynamics II

Consider next an agent watching a moving human hand generated by some simple physics-based computational model, as we described in^9^—see Figure 8.

We represent the retina image by *I* = *I*_*θ*_ ∈ ℝ^*Y*^, with *Y* the number of “pixels” in the retina (e.g., *Y* = 10^6^, the approximate number of fibers in the optic nerve^100^) and *θ* ∈ ℝ^*M*^ a small set of parameters reflecting the state of about *M* = 10^2^ ligaments and tendons (as may be generated by a realistic animation,^101^ for example). Although the space of all possible images has a large dimensionality *Y*, the space of all hand-generated images has a much smaller dimensionality *M ≪ Y*. More precisely, the points in image space corresponding to hand images lie in an *M* -dimensional manifold embedded into a *Y* -dimensional box.

The agent is equipped with a brain consisting of a large network of neurons, and the brain state specified by *x* ∈ ℝ^*X*^ (*X* is very large, there are about 1.6 × 10^10^ neurons and about 1.5 ×10^14^ synapses in the human cerebral cortex^102,103^). The dimensions of the brain, pixel, and model space are drastically different. There are many more neurons (or synapses) than data points (pixels) and very few model parameters (hand shape and orientation parameters), *M ≪ Y ≪ X*.

The generative nature of images can be understood as the action of a continuous group 𝒢 on the parameter set of the hand model and, therefore, of the images, *γ* · *I*_*θ*_ = *I*_*γ*·*θ*_. This equation states that the action of the group on image space can be understood as its action on hand parameter space. The set of all possible hand images (the hand manifold) is invariant under the action of 𝒢. This set is also equivalent to the semantic notion or the algorithmic pattern of “hand.”

We return to Equation 3.1, which becomes 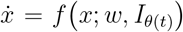, where we have expressed the model inputs as *η*(*t*) = *I*_*θ*(*t*)_ (the image may be preprocessed in some way before its input to the network—e.g., the agent may be tracking only some features of the image, not trying to match every pixel in the image —, but this is not important here).

Unless *θ* is fixed, our equation is non-autonomous. Even so, the range of possible inputs is now greatly constrained by their simple generative nature. This facilitates the agent’s task, which is modeling and (approximately) matching these inputs. We represent this by the *world-tracking constraint* : there is a projection from neural states to inputs—the neural state variable *x* carries a representation of the inputs via some projection operator (a variable selector or *projector* in practice) *p*(*x*). Putting together this with the neural dynamics equation, we have the *world-tracking neurodynamics equations (WTNE)*

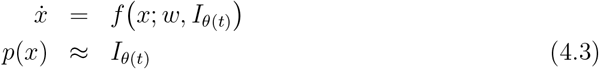

with the inputs *I*_*θ*(*t*)_ produced by the generative model and some parameter dynamics (see Figure 7).

**Figure 7:**
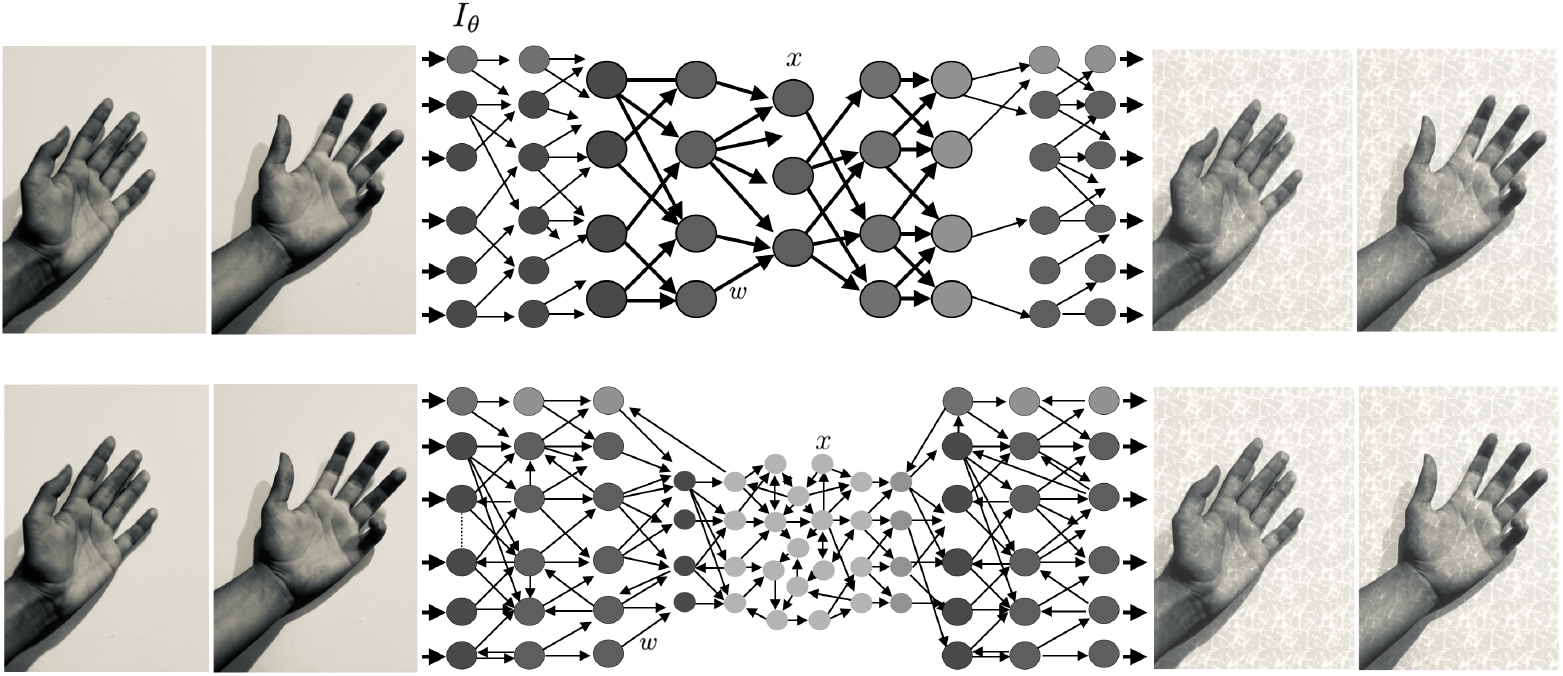
World-tracking neurodynamics. Tracking the world, represented here by a set of frames of a moving hand, is essentially the task of a compressive autoencoder, which can be described by Equation 4.3. The top panel displays a feedforward autoencoder, while the bottom provides a more general recurrent (RNN) autoencoder (connections going backward are added to highlight the potential use of predictive coding). Both realizations display an algorithmic information bottleneck (latent space), where input data is mapped to generative model parameter space.

**Figure 8:**
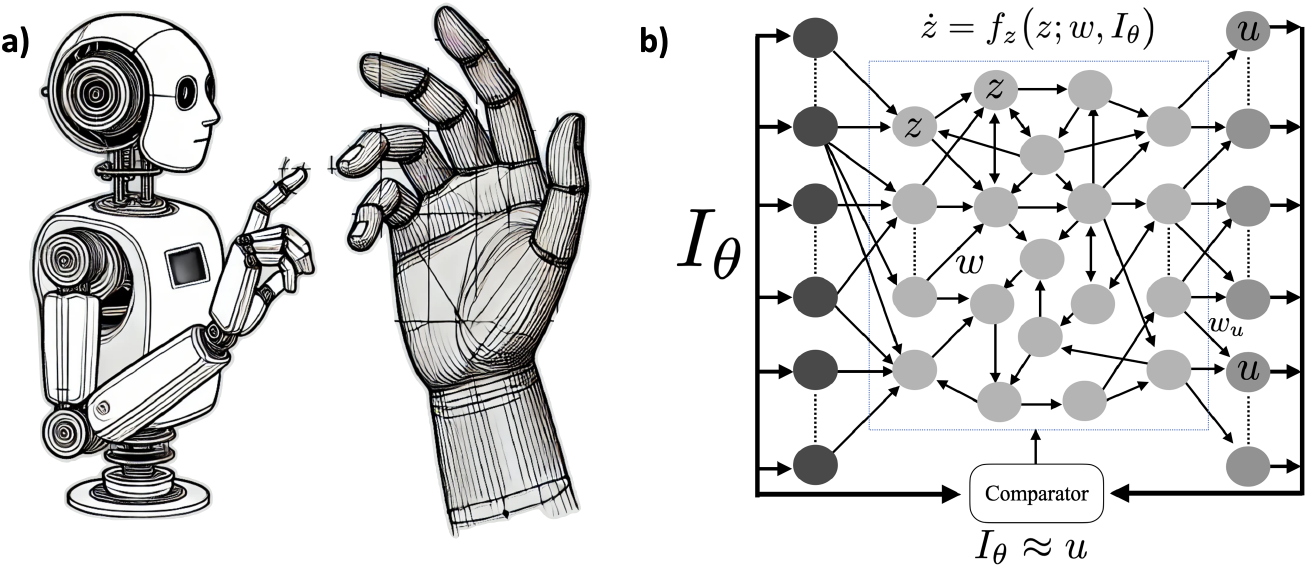
Agent tracking a computer-generated moving hand. a) An agent is observing and tracking a moving human hand generated by some simple physics-based computational model (such as Blender^101^). b) World tracking neural network, where the input is compared to the read out—see Equation 4.3. We don’t detail here how the Comparator is implemented in the network or how its output is used to regulate the network—see Section 4.4 for a more in-depth discussion. (The image in a) is AI-generated).

This a *differential algebraic equation* (it consists both of differential equations and algebraic constraints). It will not have a solution for an arbitrary set of fixed weights *w*: once the input is fixed, the system dynamics and, therefore, *p*(*x*) are determined, which will not, in general, match the inputs.

In fact, even in the case of a static input, the only way to satisfy the constraint is if the constraint is a conserved quantity in disguise,

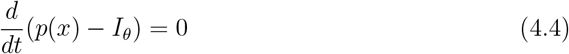

We may relax the problem by asking for approximate tracking, as indicated by the “≈” symbol in the second equation. This may be implemented by replacing the constraint with an equation of the form ‖*I*_*θ*(*t*)_ − *p*(*x*)‖^2^ < *ϵ*. We may also relax the problem by asking for approximate tracking after transient behavior (the constraint is then seen as a Lyapunov function of the ODE, see 4.4).

Another way to view Equations 4.3 is as an *autoencoder* of the input function, i.e., with the input *I*_*θ*_(*t*) approximately equal to the output *p*(*x*), as outlined in Figure 7. The network implementation is recurrent and reminiscent of architectures in Reservoir Computing,^104–106^ but it can also represent a feedforward system such as a variational autoencoder^107^—or something in between, as in Figure 7. In this machine-learning context, we rewrite the WTNE autoencoder equation as

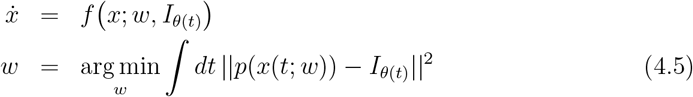

where the dataset used for training is the generated image space discussed above.

In what follows, we assume the problem has been solved, with the *w* parameters adjusted to provide such a solution (through some training process). We will also work with Equation 4.3 rather than Equation 4.5 for the sake of simplicity, although we should keep in mind that, in a real scenario, only approximate solutions will be possible.

To represent explicitly the role of Comparator, we can provide an alternative formulation in dynamical systems’ terms with more details on the world-tracking equations, where the Comparator computes the error between the model and the data (see Figure 2) and feeds it back to adjust the dynamics so that the constraint is met after a transient period. We describe the dynamics and the approximate constraint in the general WTNE (Equation 4.3) using the notion of the Lyapunov function. Lyapunov stability is a method for analyzing the behavior of dynamical systems near an equilibrium. It uses a Lyapunov function, which acts like an “energy” measure. If this function is non-increasing over time, the system is stable: small disturbances won’t cause it to diverge far from equilibrium. When the Lyapunov function strictly decreases, the system converges to a stable point or a minimum. However, if the function remains constant along certain trajectories, the system may not settle at a point but instead exhibit periodic or orbiting behavior. With this at hand, we define the dynamic world-tracking problem.

**World tracking problem formalization:** Agent dynamics (i.e., the equation structure, weights, etc.) need to satisfy the equivariant equations

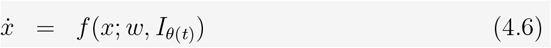

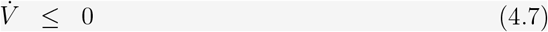

where *V* (*t*) is a Lyapunov function capturing the role of the Comparator,

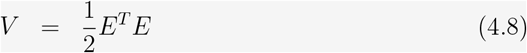

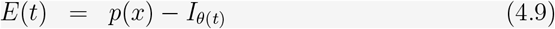

Here *x* represents the state variables of the agent system, *w* denotes the system parameters or weights, *I* _*θ* (*t*)_ is the Lie-generated time-varying input from the environment, parameterized by *θ* (*t*), *p*(*x*) is the model’s output (readout) based on the current state *x*, and *E*(*t*) is the error computed by the Comparator. The goal is to achieve world-tracking after transient behavior.

As usual, these equations are to be equivariant under the action of the Lie group. The Lie group control structure of time-varying input plays a key role in making the problem feasible by setting constraints on the equation parameters, including the weights *w*.

### 4.3 The fixed input problem

The projection operator *g* in Equations 4.3 restricts the dynamics of the state *x* through dim(*I*)=*Y* time-dependent constraint equations. The conditions set by the equations must be met with fixed weights *w* no matter what the world is doing as long as it is under the control of the generative process. In particular, if the inputs are constant (the hand is not moving or moving very slowly), we have, for any *θ*,

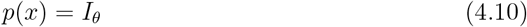

This implies that the dynamics are characterized by a large set of conserved quantities *p*_*n*_(*x*) = *C*_*n*_, *n* = 1, …, *Y* (which may not all be independent, of course). Since *p* is a variable selector, this means that some of the *x* elements will be constant (the network readout layer). The trajectories in phase space will, therefore, lie in a reduced manifold defined by the coordinates *p*(*x*) = *C*.

The existence of conserved quantities (constraints) implies the ODE has symmetry associated with them (see Appendix A.6). This implies the parameters *w* will satisfy some relations. Another way of seeing this is that the only possible way of having a solution to Equation 4.3 is if the world-tracking constraint is compatible with the conserved quantities in the ODE—not an independent new constraint, which would force the dynamics to be a single point. Expressed in (local) canonical coordinates, the world-tracking constraint must be a function of the cyclic (conserved) canonical variables. The world tracking condition requires that the system encode some conservation laws, which translate into local canonical variables where the variables corresponding to the conserved quantities are constant, i.e., 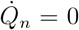. This subspace constitutes a center manifold with zero eigenvalues. This means that the generator of world inputs must be part of the symmetry group of the ODE system. Finally, as we saw, approximate symmetries can be achieved in steady state.

Finally, Equation 4.10 has to hold for any value of *θ*. Since this constraint is associated with the dynamics, 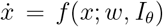, this imposes conditions on the *w* structure of the equations, as we discuss next.

#### Covariance of the equations

To analyze the general case where the network is capable of tracking *any* fixed image generated by the group, we now split the variables *x* into two blocks, *x* = (*z, u*), where *u* are the variables involved in the world-tracking constraint (see Figure 8). With some abuse of notation, since *u* = *I*_*θ*_ is constant 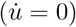, we can write the generic fixed image world-tracking equations without explicit reference to *u* as

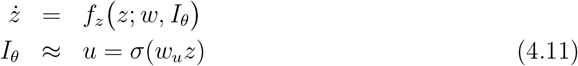

where for simplicity, we assume a feedforward read-out layer with an element nonlinearity, with *p*(*x*) = *σ*(*w*_*u*_*z*) = *u* (recall that *p* is a variable selector operator selecting *u*, a small subset of ∼ 10^6^ *x* variables). We use the approximate symbol in the second equation to allow for transients and approximate matching.

We assume the system is symmetric, i.e., we assume that, given the input *I*_*θ*_, *z* is a solution, then so is *γ* · *z* with input *γ* · *I*_*θ*_, that is,

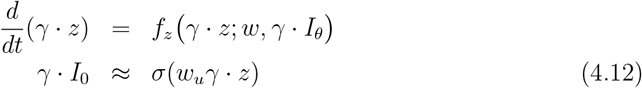

This is simply a consequence of the input tracking constraints and the Lie-generative nature of the world model. The agent is able, by definition, to track any version of the hand images (recognizing them as “hands”).

The action of the Lie group on *z* may be nonlinear, but the condition applies to an arbitrary infinitesimal transformation *γ* = 1 + *ϵ*^*k*^*T*_*k*_(*z*), i.e., in the linear regime (with *T* a linear operator, see^72^ for a more formal mathematical discussion.). These operators commute with time differentiation (*T*_*k*_(*z*) act on configuration space), hence

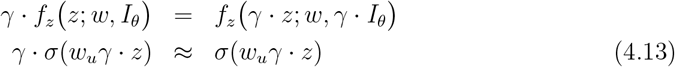

for an infinitesimal transformation. Expanding *γ* = 1 + *ϵ*^*k*^*T*_*k*_(*z*), we can write a pair of such equations for each generator *T*^*k*^ (linear operator) since *ϵ* is arbitrary and the generators linearly independent.

Covariance under the Lie group transformations, Equation 4.13, imposes specific conditions on the weights *w*. These conditions ensure that the dynamics *ż* = *f*_*z*_(*z*; *w, I*_*θ*_) are consistent with the symmetry properties of the system and the desired conservation laws (world tracking constraints).

As a consequence of the world tracking constraints associated with inputs from a generative model, we see that the number of conservation laws or constraints is proportional to the number of group generators. This is because each generator of the Lie group introduces constraints through its action on the output variables *u*. A group with a large number of generators will impose tighter constraints (and more conservation laws) on the possible *w* space.

### 4.4 Tracking time-varying inputs

Here, we extend the analysis to the situation where the inputs change in time but at a long time scale compared to neural dynamics to allow for transient phenomena to complete. As the input image moves slowly through changes in the *M* -dimensional parameters *θ* in the generative model, the reduced manifold of trajectories will also shift. Through this process, the manifold will increase its dimensionality, but it can only acquire a maximum of *M* extra dimensions. Because the world model is assumed to be simple, i.e., with *M* small, even in the case where the parameters *θ* are dynamic, the invariant manifold of the tracking system will remain dimensionally constrained.

The fact that images are generated by a simple model using a few parameters compared to the dimensionality of image space is crucial. In the example above, we analyzed the case where the image does not vary in time, i.e., a static input. If, on the contrary, the image is fully dynamic (each pixel varying in time independently of the others), there would be no reduction of the dimension of the trajectory space. There would be as many constraints as pixels, but they would be time-dependent, and possibly, no satisfactory weights *w* would be found in such a case. It is the intrinsic low-dimensionality of data generation that allows almost all the constraint equations to independently contribute conserved quantities: the image *I*_*θ*_ lies in a small dimensional manifold, and the operator *g* must project *x* into it.

The world-tracking equations are expected to hold in an approximate way and only after a transient time. As in the simple example in Section A.6.3, it may be the case that the world-tracking constraints are realized as attractors in the dynamics. In the case of a dynamics input generated by dynamical parameters in the generative model, we can expect to see an approximate match of the output to the input only after a transient period of time.

So far, we have not discussed *how* the agent system can track changing inputs. We now provide a potential solution to the problem in the form of a specific structure for Equation 4.6 assuming that only the error is used by the network using proportional feedback,^108^

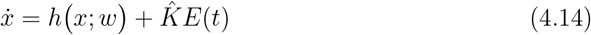

where 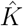 is a linear feedback gain operator that determines the influence of the error on the system’s dynamics.

The time derivative of *V* is 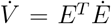. Substituting the dynamics of *Ė* from Equation (4.14):

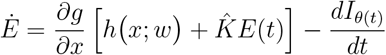

To ensure 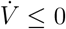, the feedback gain 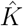 must be designed such that

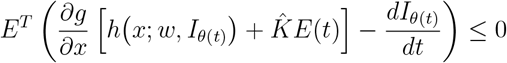

Proper selection of 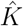 will ensure that *V* (*E*) is non-increasing, thereby driving *E*(*t*) towards zero asymptotically. The choice of this operator will reflect the Lie group structure of the system since these equations need to be equivariant. We leave this analysis for further work.

Finally, these equations correspond to the fast-time equations in Ruffini et al., 2024.^29^ The Comparator error data stream *E*(*t*) will feed the slow-time *connectodynamics* equations to govern the slow change of the structural parameters *w*, 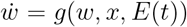, with *g*(*w, x*, 0) = 0.

In summary, this section extends the previous analysis of static inputs to the case of slowly varying signals mediated by the Lie group action, showing that even when the environment undergoes incremental changes, the low-dimensional structure of the generative model and equivariance can provide a tractable path for world-tracking with an appropriate selection of weights and gain operator. Thus, the same principles that apply to static generative scenarios—namely, leveraging low-dimensional structure and compositional transformations—remain central for handling time-varying inputs. This completes the foundations laid in earlier sections, formalizing how compositional and equivariant mechanisms enable agents to track and compress real-world complexity in both static and slowly evolving contexts.

### 4.5 Coarse-graining, hierarchical constraints, and manifolds

An important idea in the algorithmic agent framework is that all cognitive processes— modeling, planning, and objective function evaluation—are inherently hierarchical (compositional).^9^ This means decision-making and goal-setting occur in layers, ranging from simple actions to complex behaviors. This hierarchical organization naturally finds its origin in the multi-scale structure of real-world data enabling effective modeling and decisionmaking. To ensure survival with limited resources, the agent must employ *coarse-graining* to transform complex, uncompressible data into a simpler, compressed form while retaining a non-trivial structure.^11^ Techniques like spatiotemporal averaging, spectral decomposition, compressive sensing, and dimensionality reduction achieve this by reorganizing data at various levels of abstraction. This perspective aligns with the concept of the brain processing information through *hierarchical coarse-graining*, as supported by research in the visual^109,110^ and auditory systems.^111^ A similar approach is utilized by deep neural network architectures, including convolutional neural networks.^68,73^

At each processing level, details are aggregated and abstracted to form higher-level representations. World-tracking at different levels corresponds to different scales of coarse-graining, progressively reducing the complexity of the system while preserving essential features. Since lower-level constraints (e.g., “it’s furry”) must be compatible with higher-level constraints (“it’s a cat”), there is a natural nested structure of constraints and corresponding manifolds (which also induces an inclusion relation among their tangent spaces).World tracking can be achieved through a hierarchy of constraints {𝒞_*i*_}^*k*^, each corresponding to a specific coarse-graining level. The constraints are nested and must be compatible, meaning each lower-level constraint operates within the subset of the state space defined by the higher-level constraints.

We can formalize this concept in a simplified manner (reality is undoubtedly more complex, involving parallel threads of constraints, for example) through a sequence of *coarsegraining operators* 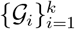, where each operator 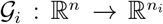 maps the fine-grained state vector **x** to a coarse-grained state vector **y**_*i*_, with **y**_*i*_ = 𝒢_*i*_(**x**) (*n*_*i*_ *< n*). Each **y**_*i*_ represents the system at a specific level of abstraction, capturing variables relevant at that scale. For each level *i*, there is a corresponding *hierarchical constraint* 𝒞_*i*_ on **y**_*i*_, expressed as 𝒞_*i*_(**y**_*i*_) = 0. Constraints reflect the agent’s world model at different levels, ensuring that the dynamics of the coarse-grained variables align with expected behaviors.

Lower-level constraints must be *compatible* with higher-level constraints. This means that the solution set of a lower-level constraint 𝒞_*i*+1_ is a subset of the solution set of the higher-level constraint 𝒞_*i*_. Formally, {**x** ∈ ℝ^*n*^ | 𝒞_*i*+1_(𝒢_*i*+1_(**x**)) = 0} ⊆ {**x** ∈ ℝ^*n*^ | 𝒞_*i*_(𝒢_*i*_(**x**)) = 0}. Nesting ensures that as we move to lower levels (higher indices), the constraints become more specific, refining the state space within the context defined by higher-level constraints.

As an example, consider a visual perception task with a high-level constraint 𝒞_1_: the agent recognizes that the object is a *cat* —see Figure 9. This constraint reduces the state space to the manifold ℳ_1_ of all possible cats, which is still high-dimensional. To this, we can add a lower-level constraint 𝒞_2_: the agent further discerns that the cat is *white with blue eyes*. This constraint reduces the state space to a submanifold ℳ_2_ within ℳ_1_, containing only white cats with blue eyes. Here, the lower-level constraint 𝒞_2_ is compatible with the higher-level constraint 𝒞_1_, as all white cats with blue eyes are indeed cats, ℳ_2_ ⊆ ℳ_1_. Thus, higher-level constraints define broad categories or contexts (e.g., recognizing an object as a cat), while lower-level constraints provide additional specificity (e.g., identifying the cat’s color, hair type, etc). The nested structure ensures that all lower-level processing is coherent with the higher-level understanding, leading to efficient and accurate modeling of the world. The hierarchical framework of constraints is related to compositional Lie groups characterizing the invariant properties of symmetries of the equations. In particular, the hierarchical constraints (𝒞_*i*_) on agent dynamics correspond to conserved quantities associated with symmetries of the ODEs represented by a hierarchical Lie group structure.

**Figure 9.**
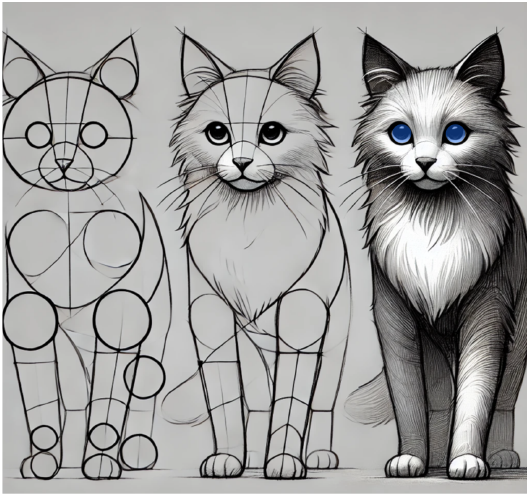
Hierarchical modeling: step-by-step drawing of a cat demonstrating hierarchical constraints: starting with the basic form of a cat (“it’s a cat”), refining it with more specific features, and finally adding details like “has white fur and blue eyes,” progressively narrowing the state space to match more specific characteristics. (Image is partially AI-generated.)

By imposing these hierarchical, compatible constraints, the original high-dimensional phase space ℝ^*n*^ is reduced in a sequence of *nested manifolds*, forming what we may call a *hierarchical or compositional manifold*,

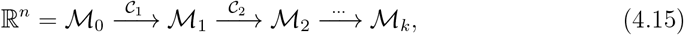

where each manifold is defined recursively as

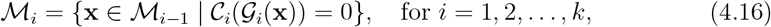

with ℳ_0_ = ℝ^*n*^. Each manifold ℳ_*i*_ satisfies all constraints up to level *i*, ensuring that the agent’s internal states remain consistent with both high-level goals and fine-grained observations. This recursive structure ensures that ℳ_*i*_ ⊆ ℳ_*i*−1_,, reflecting the increased specificity of constraints at lower levels (this also induces an inclusion relation among their tangent spaces).

The sequence of compositional compatible constraints leads, in turn, to a sequence of nested manifolds, each representing the state space under the accumulated constraints up to that level. The agent’s dynamics are thus confined to a reduced hierarchical manifold, ensuring that its internal states remain consistent with both high-level goals and fine-grained observations. This mirrors the hierarchical organization of the brain or deep artificial networks, where a nested structure of coarse-grained models synthesizes a multilevel emergent view of the world.

## 5 Discussion

In this paper, we first assumed that agent inputs from the world are generated by simple, hierarchical rules (i.e., that the world is inherently simple in algorithmic terms), i.e., by a hierarchical (compositional) Lie generative model, and that the agent can track its inputs, i.e., encodes an internal generative model to approximately match data, perhaps after transients. From this, we showed that

1. As a dynamical system, the successful agent displays conserved quantities— with dynamics in a reduced hierarchical manifold—corresponding to the world-tracking constraints generated by a Lie group. For dynamical inputs, this stability is achieved after initial transients, captured by a Lyapunov function (see Section 4.4).
2. The constraints force *structural* symmetries in the dynamical system constitutive equations that meet them: the agent carries the symmetries of the world generative model. This is encoded in the structural elements of the dynamical system (the *w*s in our formulation).

World-tracking conditions are satisfied by specialized, symmetric networks, as the governing equations must maintain (approximately) conserved quantities that are compatible with the required outputs. Conversely, constraint (symmetry) breaking, as detected by the agent in the Comparator (Figure 2), allows the agent to identify anomalies or novel patterns, prompting updates to its internal models and engaging in learning and adaptation.

If the inputs are static, dynamics will lie in a reduced manifold, as determined by the constraints of world tracking. If the inputs vary slowly—following the rules of a generative world model—the dimension of the reduced manifold will increase, but only slightly compared to the embedding neural dimension. If inputs violate the world model, on the other hand, tracking will fail, and the dynamics may no longer be constrained.

Our analysis thus links the *manifold hypothesis* with world tracking and symmetry principles. Empirical observations of dynamics in natural systems indicate that data trajectories typically lie in a manifold of dimensionality much lower than ℝ^*X*^ (the invariant manifold), which we also loosely identify with the *latent space* when using autoencoders to compress neural data. The manifold hypothesis posits that high-dimensional data, such as world data collected from sensory systems (e.g., images) or neuroimaging, can be compressed into a reduced number of parameters (latent space^112^) due to the presence of a low-dimensional manifold within the high-dimensional phase space.^63,65,113,114^ Such manifolds can take various forms, such as attractors and unstable manifolds.

Because the data-generating processes we consider are inherently low-dimensional, our analysis provides a link with the *manifold hypothesis* in agent-generated data: large-scale data sets (be they high-dimensional images or neural recordings) often occupy manifolds of significantly lower dimension than the raw data space would suggest.^63,65,113,114^ Here, we see that if data truly arise from a compositional world model, then an agent tracking those data automatically inherits such a manifold structure in its own state space, effectively compressing high-dimensional inputs into a smaller number of effective degrees of freedom.

Finally, this compositional view also suggests a *hierarchical* structure of manifolds when world data are generated at multiple, nested scales of abstraction, yielding the notion of a *hierarchical manifold*. This hierarchical perspective emphasizes how multiple levels of coarse-grained, low-dimensional constraints can collectively shape both world data and agent dynamics, providing different spatiotemporal scales of shared algorithmic information (see, e.g.,^115^ for related concepts in the Shannon information theory context). From this vantage, compositional world data and the act of tracking them together naturally realize the manifold hypothesis in both observational data and in an agent’s neural (or computational) trajectories.

A reduced hierarchical data manifold embodies the inherent simplicity of the underlying generative models that produce the observed data, as well as the effectiveness of a world-tracking system in learning these models. This observation supports the **manifold hypothesis** as it manifests in world-tracking systems.

### 5.1 Symmetry, groups and algorithmic complexity

Does group action encompass the general notion of compression — finding short programs (i.e., with low Kolmogorov Complexity)? Under what conditions does the existence of a highly compressive generative model imply the presence of underlying symmetries described by a group (or generalization of)? Although rigorously answering this will require bridging abstract algebra and algorithmic information theory, the intuition is straightforward: a compressive (short) program capable of generating a large dataset will involve repeating an operation (recursion) or a sequence of operations (compositionality) multiple times.

This suggests that in a compressive model, the recursive action of a Lie group in configuration/latent space is represented economically in the construction of the generative function. In particular, if the image-generating function is a huge lookup table, there will be no compression despite the reduction of dimensionality from image to latent space. The *recursive* property of Lie groups, where complex elements are constructed through repeated applications of simple algebraic operations, mirrors foundational concepts in algorithmic information theory and recursive function theory. *Compositionality* in this context refers to the combined action of different generators.

The presence of a Lie group structure in a generative model significantly influences its algorithmic complexity. If one can navigate a configuration space with a Lie group, any function on that space—including a generative model—can be manipulated using the Lie group’s generators. This approach can be highly efficient. By generating a single instance within our model, we can apply the Lie group’s transformations in small, discrete steps to produce a diverse set of outputs.

As we saw, this process is inherently *recursive* and *compositional*, hence enabling a compressive representation. The program only needs to encode the initial instance of the object and the transformation rules derived from the Lie group’s generators, thus enabling the generation of a vast array of images through a sequence of incremental, systematic transformations.

Finally, we recall that the *compositional sparsity* of the underlying target function (i.e., the world)—the task to be learned—is the key principle behind the success of machine learning, particularly deep learning architectures such as CNNs or transformers.^87,116^ Compositional sparsity, the idea that a function can be computed through the composition of functions, each with a constituent function of low dimensionality,^117^ is equivalent in algorithmic terms to efficient computability, that is, computability in polynomial time.^116^ Thus, algorithmic and computational complexity notions may converge on the role of Lie action in generative (compressive) models.

In summary, we highlighted the relationship between algorithmic complexity and Lie groups, which stems from compressive models aiming to capture and recreate complex data structures through *recursion and compositionality* and the natural connection of these and Lie group action.

### 5.2 Connections with empirical observations

Symmetry is an example of an algorithmic feature that systems that compute the same function must share, regardless of their specific implementation. This implies that natural agents such as human brains implement symmetries inherent in world data within their architecture and, in turn, generate data with conserved quantities. This translates into reduced dimensionality dynamics in invariant manifolds and may explain complexity features observed in neuroimaging data across brain states and neurological or psychiatric disorders. Techniques like topological data analysis may help uncover further correlates of structured experience with applications in consciousness research in natural and artificial systems.

The fundamental predictions from studying the role of symmetry in agent dynamics pertain to the structural constraints of its constitutive equations and to the emergence of reduced manifolds, reflecting the structure of compositional data the agent is tracking.

The simplicity associated with a generative model may manifest in different ways. For instance, an efficient autoencoder of neural data from a brain exposed to time-varying world inputs could regenerate the dynamics using a latent parameter space whose dimension approximately matches the dimension of the world-generative model configuration space.

Furthermore, the Comparator mechanism enables the emergence of Mutual Algorithmic Information (MAI) between the agent and the world, which KT postulates.^8,9^ By aligning internal models with external data, the Comparator highlights how constraints in world-tracking impact the structure of reduced, invariant manifolds in neural dynamics. These manifold structures, in turn, are expected to vary with brain states and environmental conditions, influencing neuroimaging data across different states of consciousness (awake, REM, NREM),^112^ under anesthesia, minimally conscious (MCS) or unresponsive wakeful state (UWS), Locked-In Syndrome (LIS), epilepsy or Alzheimer’s disease (AD),^118^ which should display different latent space features, including dimensionality (compressibility, complexity), or exhibit task dependence (e.g., eyes open or closed). This can be attributed, at least in part, to the world-tracking features associated with brain state or task and motivates the hypothesis that structured experience is associated with the successful match of data and model at the Comparator.^8–10^

Constrained dynamics may also provide the context for phenomena such as increased EEG slow-frequency power in the eyes-closed state (and reduced complexity^119^), anesthesia, or completely locked-in syndrome (CLIS, where patients shift toward slower frequencies and lower signal complexity),^120,121^ all of which are characterized by clamping down (setting to a null fixed value) world inputs. This may either reflect suppression of Comparator outputs or the active tracking of a predicted static input. On the other extreme, the perspective of constrained dynamics may explain how psychedelics, which interfere with sensory processing, disrupt the Comparator and modeling process (Equations 4.6–4.9), with network dynamics departing from the normative invariant manifold. Such a departure manifests as an increase in complexity and reduction in alpha power.^42,43,122–127^

For data analysis, approaches such as deep learning autoencoders are well-suited for uncovering latent spaces associated with brain dynamics. Complementing this, topological data analysis can reveal structural features of reduced manifolds. These techniques could provide new insights into how different states compress or expand neural dynamics, providing insights into underlying brain symmetries and their breakdown in pathology. Complexity measures, like the Lempel-Ziv algorithm, can help quantify the intrinsic dimensionality of these manifolds, revealing symmetry-related information processing characteristics.^17,128^

Finally, while we have focused here on how compositional data-tracking gives rise to constrained dynamics, the same logic applies to the generation of *compositional outputs* by the agent, such as motor actions. Recent research has uncovered low-dimensional neural manifolds that capture significant neural variability across various brain regions and experimental conditions.^129–138^ These manifolds, defined by correlated neural activity patterns of neural activity —referred to as “neural modes,”— are proposed to generate motor behavior through their time-dependent activation.^129,133^

This research will benefit from a similar analysis of artificial systems, as discussed next.

### 5.3 Connections with neurophenomenology

The emergence of reduced manifolds governed by Lie pseudogroups, as described in the context of agent-world interactions, offers a link to neurophenomenology. When an agent tracks compositional data from the environment, its internal dynamics are constrained to invariant manifolds reflecting the structure of the data. In this framework, these reduced manifolds shape the qualia—the fundamental units of structured experience associated with the agent’s perception of reality.

Neurophenomenology, which seeks to bridge first-person experiential phenomena with third-person neural dynamics, can provide methods to validate this perspective. Structured experiences can be viewed as arising from the agent’s successful alignment of internal models with world data, as mediated by the Comparator mechanism. This suggests combining the characterization of brain states by the geometry and topology of the underlying neural manifolds, which reflect the agent’s interaction with its environment together with phenomenology methods^139^ for the characterization of the structure of experience— see Figure 1.

The integration of psychophysics, phenomenology, and AI methods,^140^ such as natural language processing (NLP),^141–143^ offers a powerful approach to characterizing the structure of experience. Psychophysics provides quantitative tools to link physical stimuli with subjective perception, while phenomenology delves into the qualitative aspects of experience, emphasizing first-person reports and the lived reality of consciousness. AI methods like NLP can bridge these domains by analyzing and modeling linguistic data, enabling the systematic study of subjective descriptions. This combination allows for a multi-dimensional exploration of structured experience, where psychophysics anchors the investigation in measurable variables, phenomenology enriches it with introspective depth, and AI scales and systematizes the analysis, revealing patterns and latent structures in experiential data. Together, these tools provide a framework for advancing our understanding of the dynamics and features of structured experience.

### 5.4 Discovering structure

In this paper, we have discussed the implications of using compressive models of generative world data, but we have not touched on the fundamental issue of how these models are discovered. A potential insight in this direction is that recursion and compositionality are the key features of algorithmics.^7^

Recent research indicates that deep neural networks inherently favor low-complexity, compositional data structures, enabling them to identify and represent underlying symmetries without explicit prior knowledge. This intrinsic bias allows for effective generalization across various domains, supporting the unification of diverse tasks within a single learning algorithm.^144,145^ The hierarchical and compositional nature of deep networks facilitates efficient approximation of complex functions, mitigating the curse of dimensionality and enhancing performance on tasks with compositional structures.^87,88,116,146^ These characteristics underscore the adaptability and efficiency of deep learning models in processing structured, real-world data.

Neural networks can learn to detect group structure (Lie algebras) from training data, and extracting invariances remains an active research area. Moskalev et al.^78^ propose a two-step procedure: (i) train a network on a task requiring invariance, and (ii) extract the Lie algebra that underlies this learned symmetry. Their Lie Group Generators (LieGG) method builds a polarization matrix from the network’s input-output relationship, applies singular value decomposition, and identifies near-zero singular values that correspond to the basis of the learned Lie algebra. For instance, on rotation MNIST, LieGG recovers the rotation symmetry and uses metrics like “symmetry variance” and “symmetry bias” to assess how thoroughly the network has internalized this invariance.

Recent advancements have introduced additional methods for discovering symmetries and invariances in data using neural networks. For example, LieSD (Lie Symmetry Discovery) identifies continuous symmetries by analyzing the gradients and outputs of trained neural networks, demonstrating its utility in tasks like the two-body problem and top quark tagging.^147^ Similarly, LaLiGAN (Latent LieGAN) learns a latent space representation where nonlinear symmetries become linear, enabling the discovery of intrinsic symmetries in high-dimensional systems.^148^ Another approach, Lie Algebra Convolutional Networks (L-conv), uses Lie algebras instead of groups to construct group-equivariant architectures, facilitating symmetry discovery and connection to physical principles like conservation laws.^149^ These methods extend the capabilities of LieGG, enabling the exploration of both geometric and more abstract symmetries in neural networks.

Another perspective is exploring the topology of Lie groups^150^ (and, more generally, Lie pseudogroups), which provides an essential framework for understanding the ways in which agents track and encode structured data in their internal dynamics. In particular, based on topological data analysis, properties such as connectedness and the presence of non-trivial loops or higher-dimensional “holes” (often characterized by Betti numbers) can inform how a neural system might transition among configurations or maintain stable invariants in response to environmental inputs. Within neurophenomenology, such topological features of the agent’s internal manifold—mirroring the underlying compositional actions of a Lie group—can shape how perceptual states coalesce into coherent experiences.

### 5.5 Applications in AI and neurosynthetic agent design

In *artificial intelligence* (AI) and *computational neuroscience*, symmetry constraints are pivotal in guiding model design by imposing structural limitations that reflect the symmetries inherent in World interaction tasks. These symmetry principles can be incorporated as regularization terms to reduce parameter degeneracy or structurally (e.g., convolutional networks), enhancing model robustness and efficiency. From the earliest developments of deep networks^68,73,74,151^ to current advancements, symmetry-driven approaches continue to shape and fuel research into more adaptable and efficient models.^69,71,78,83,152^

Whole-brain computational models and digital twins^153^ are *neurosynthetic* neural networks characterized by a very large number of parameters that are difficult to adjust given the limited available neuroimaging data. Just as with artificial neural networks, enforcing symmetry and conserved quantities with low dimensionality, we can reduce the parameter space’s degeneracy when fitting models to data.^154^ Additionally, when parameters are known, the system’s dynamics can be explored through its symmetry properties. Analyzing the number, dimension, structure, and stability of the system’s attractors (using eigenvalue and bifurcation analysis) provides insight into its long-term behavior^1,95^ and computational capabilities, including multistability, metastability, and criticality.^34, 40, 155, 156^

For instance, whole-brain models need to reflect the spatial-temporal symmetries and sensory-motor contingencies inherent in the data they process.^157^ This includes fundamental transformations like translations and rotations. Such considerations will become increasingly important as whole-brain models are developed to interact with external inputs and generate outputs, whether in classification tasks or motor actions.

An important part of this program is to characterize more general world symmetries and translate them into generative world models and Lie groups. An approach to attack this may be the use of deep learning models that can produce the appropriate latent spaces or derive the Lie group generators from networks that have been trained on world data.^70,72,78^ The discovery of these attractors can be used to explore the mutual algorithmic information between data and dynamical attractors in the artificial system (or in the human brain).

Finally, symmetry considerations can serve as a regularization principle, instrumentalizing a bias towards simplicity. By demanding that models remain as symmetric as possible while fitting empirical data, we implement a form of Occam’s razor, promoting simpler, more generalizable models. This principle is particularly relevant for brain-inspired networks, where preserving computational simplicity can improve interpretability and robustness.^1^

## 6 Conclusion

In past work, we put forth the idea that because agents possess and run models of the World, there is significant Mutual Algorithmic Information (MAI) between the agent and the world.^8^ In this paper, this idea is made concrete in dynamical terms using the language of group theory: the agent, viewed as a dynamical system, encodes symmetries that reflect the structure of world data.

The historical trajectory of group theory—the mathematical theory of symmetry—is a testament to its fundamental role in shaping the landscape of modern mathematical and physical sciences. From its initial development by Évariste Galois and Paolo Ruffini^158–160^—who, to unveil the algebraic symmetries governing polynomial equations, created the seeds of group theory—, to the works of Sophus Lie,^1,95^ who extended these concepts into the role of continuous groups in differential equations, group theory is rich in its fruits. The subsequent epoch, marked by Emmy Noether’s transformative theorem,^66,67^ cemented the profound connection between symmetries and conservation laws, further elucidating the inherent simplicity underlying complex natural phenomena and driving the development of General Relativity and the Standard Model in physics.

These historical milestones underscore the enduring quest for understanding through the lenses of simplicity and symmetry, to which we hope this paper contributes. As we have discussed here through the model of the algorithmic agent (KT), the convergent principles of algorithmic simplicity and symmetry can yield useful insights into the nature of artificial and natural intelligence and the deep connections between mathematics and the phenomenology of structured experience.

## Acknowledgments

The author wishes to thank Professor Motohico Mulase (Distinguished Professor of Mathematics at the University of California, Davis) for useful comments on Lie group theory, pseudogroups, and moduli stacks.

## A Appendix

### A.1 Equations for a Turing pair

The general equations for a *Turing pair* —a description of the interaction of the agent and world—implemented as a network dynamical system are given by the autonomous system

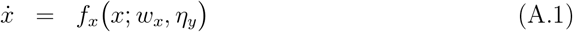

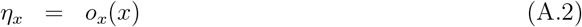

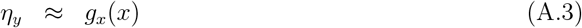

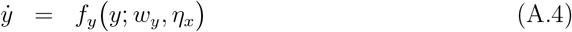

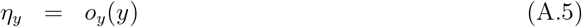

Or, explicitly providing the Comparator equation with the Lyupanov function,

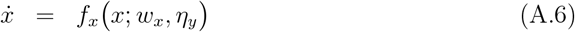

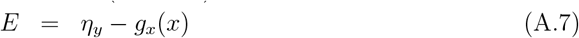

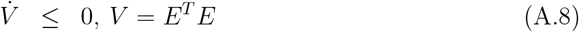

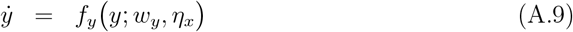

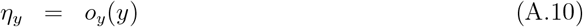

These equations express the idea that the outputs of a subsystem are the inputs of the other. They include the agent’s world-tracking condition. Here, *x* is a multidimensional variable capturing the state of the agent, and *y* captures the state of the world. The *w*s are connectivity weights in the network (or other parameters), and the *η*s represent the outputs of a subsystem, which become inputs for the other. The *o* functions are variable selectors of the subset of variables that become outputs. Finally, *g*_*x*_ is a variable selector (or, more generally, readout function) in the agent that is to match the inputs from the world (the world-tracking subsystem). The third equation, *η*_*y*_ = *g*_*x*_(*x*), or its Lyupanov version (allowing for transients), captures the world-tracking condition.

### A.2 Notes on Lie groups

#### A.2.1 Definitions

##### Definition A.1.

A **group** is a set *G* together with a group operation, usually called multiplication, such that for any two elements *g* and *h* of *G*, the product *g* · *h* is again an element of *G*. The group operation is required to satisfy the following axioms:

1. **Associativity**. If *g, h*, and *k* are elements of *G*, then

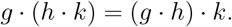
2. **Identity Element**. There is a distinguished element *e* of *G*, called the identity element, which has the property that

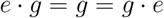

for all *g* in *G*.
3. **Inverses**. For each *g* in *G*, there is an inverse, denoted *g*^−1^, with the property

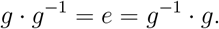

##### Definition A.2.

An *m***-dimensional manifold** is a set *M*, together with a countable collection of subsets *U*_*α*_ ⊆ *M*, called *coordinate charts*, and one-to-one functions *x*_*α*_ : *U*_*α*_ → *V*_*α*_ onto connected open subsets *V*_*α*_ ⊆ ℝ^*m*^, called *local coordinate maps*, which satisfy the following properties:

1. The coordinate charts cover *M* :

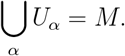
2. On the overlap of any pair of coordinate charts *U*_*α*_ ∩ *U*_*β*_, the composite map

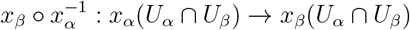

is a smooth (infinitely differentiable) function.
3. If *x* ∈ *U*_*α*_, *x*^′^ ∈ *U*_*β*_ are distinct points of *M*, then there exist open subsets *W* ⊆ *V*_*α*_, *W* ^′^ ⊆ *V*_*β*_ with *x*_*α*_(*x*) ∈ *W, x*_*β*_(*x*^′^) ∈ *W* ^′^, satisfying

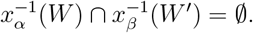

##### Definition A.3.

An *r*-parameter **Lie group** is a group *G* which also carries the structure of an *r*-dimensional smooth manifold in such a way that both the group operation

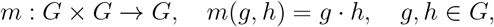

and the inversion

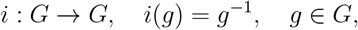

are smooth maps between manifolds.

In general, a Lie group *G* will be realized as a group of transformations of some manifold *M* with each group element *g* ∈ *G* associated with a map from *M* to itself.^1^ It is important *not to restrict* our attention solely to linear transformations. Moreover, the group may act only locally, meaning that the group transformations may not be defined for all elements of the group nor for all points on the manifold.

##### Definition A.4.

Let *M* be a smooth manifold. A local **group of transformations** acting on *M* is given by a (local) Lie group *G*, an open subset 𝒰, with

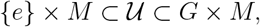

which is the domain of definition of the group action and a smooth map

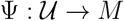

with the following properties:

1. If (*h, x*) ∈ 𝒰, (*g*, Ψ(*h, x*)) ∈ 𝒰, and also (*g* · *h, x*) ∈ 𝒰, then

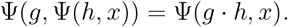
2. For all *x* ∈ *M*,

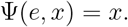
3. If (*g, x*) ∈ 𝒰, then (*g*^−1^, Ψ(*g, x*)) ∈ 𝒰 and

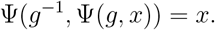

(Note that except for the assumption of the form of the domain 𝒰, part (c) follows directly from parts (a) and (b).)

For brevity, we will denote Ψ(*g, x*) by *g* · *x*, and the conditions of this definition take the simpler form:

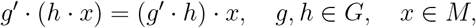

whenever both sides of this equation make sense.

##### Definition A.5.

A **Lie pseudogroup 𝒢** on a smooth manifold *M* is a collection of local diffeomorphisms *ϕ* : *U* → *V*, where *U, V* ⊆ *M* are open subsets, satisfying the following properties:

1. **Identity**: The identity map id : *U* → *U* for any open subset *U* ⊆ *M* belongs to 𝒢.
2. **Closure under composition**: If *ϕ* : *U* → *V* and *ψ* : *V* → *W* belong to 𝒢, then their composition *ψ* ∘ *ϕ* : *U* → *W* also belongs to 𝒢.
3. **Closure under inversion**: If *ϕ* : *U* → *V* belongs to 𝒢, then its inverse *ϕ*^−1^ : *V* → *U* also belongs to 𝒢.
4. **Closure under restriction**: If *ϕ* : *U* → *V* belongs to 𝒢 and *U* ^′^ ⊆ *U, V* ^′^ = *ϕ*(*U* ^′^), then the restriction 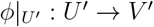 also belongs to 𝒢.
5. **Lie structure**: The local diffeomorphisms in 𝒢 are solutions of a system of finite-order partial differential equations defined on *M*, ensuring 𝒢 has the structure of a smooth (Lie) manifold.

#### A.2.2 Important theorems

This theorem shows that smoothness implies that the behavior of the group is already specified in any small patch containing the identity. In a sense, Lie groups are “cyclical”: all the elements can be obtained by repeated action of a subset.

##### Proposition 1.24.

^**1**^ Let *G* be a connected Lie group and *U* ⊆ *G* a neighborhood of the identity. Also, let

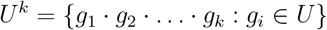

be the set of *k*-fold products of elements of *U*. Then

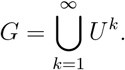

In other words, every group element *g* ∈ *G* can be written as a finite product of elements of *U*.

Furthermore, if *G* is a *connected, compact* matrix Lie group, the exponential map for *G* is subjective (covers the entire group).^75^

Related to this, one can show that there is a one-to-one relationship between a local Lie group and the Lie algebra that describes its behavior near the identity (v. Theorem 1.54 in^1^). The latter can be fully described by a set of constants (the structure constants of the Lie algebra). Thus, finite Lie groups are intrinsically simple objects.

We next address the issue of local transitivity.

##### Proposition 1 (Local Transitivity).

Let *C* be the configuration space manifold. For any point *p* ∈ *C*, there exists a neighborhood *U*_*p*_ around *p* and a Lie group *G*_*p*_ such that *G*_*p*_ acts transitively on *U*_*p*_ and transforms the local configurations of images within this neighborhood.

*Proof*. We need to show the existence of a Lie group that acts transitively on a local patch around any point *p* in *C*. At any point *p* in *C*, there exists a tangent space *T*_*p*_*C*, which captures the infinitesimal displacements around *p*. This tangent space is a vector space, and its dimension is the same as the dimension of *C*. The general linear group *GL*(*n*, ℝ), where *n* is the dimension of *T*_*p*_*C*, can act on *T*_*p*_*C* by matrix multiplication. This action is transitive since for any two vectors *v, w* in *T*_*p*_*C*, there exists a matrix *M* in *GL*(*n*, ℝ) such that *M* · *v* = *w*. By the above point, around every *p*, there exists a local action of a Lie subgroup of *GL*(*n, ℝ*) that acts transitively on a local patch *U*_*p*_ around *p* in *C*. We can take *G*_*p*_ to be this Lie subgroup. Hence, for every point *p* in *C*, we have found a Lie group *G*_*p*_ that acts transitively on a local patch *U*_*p*_ around *p*.

For the case of Lie pseudogroups, a similar theorem is available:^161^

##### Theorem A.1

(Transitivity of a Lie Pseudogroup). Let 𝒢 be a Lie pseudogroup acting on a connected smooth manifold *M*, defined by a system of partial differential equations. If G is **locally transitive** (i.e., for every point *p* ∈ *M*, there exists a neighborhood *U* ⊂ *M* where G acts transitively on *U*), then 𝒢 is **globally transitive** on *M*.

*Sketch of Proof*

1. **Local Transitivity:** By assumption, for any *p* ∈ *M*, the pseudogroup 𝒢 connects any two points within a sufficiently small neighborhood *U* ⊂ *M*.
2. **Connectivity of** *M* : Since *M* is connected, local neighborhoods *U* overlap, allowing compositions of transformations to propagate the action of 𝒢 across the entire manifold.
3. **Closure of 𝒢:** The closure properties of 𝒢 under composition and inversion ensure that transformations can be chained together to connect distant points.
4. **Conclusion:** By iteratively combining local transformations, 𝒢 acts transitively on *M*.

In the classical setting, if a finite-dimensional Lie group *G* acts *transitively* on a manifold *L*, then *L* is globally a *homogeneous space* of the form *G/H*, where *H* is the stabilizer subgroup of a chosen reference point. This immediately constrains the *dimension* of *L*, dim(*L*) = dim(*G*) − dim(*H*), and endows *L* with the same topology as the coset space *G/H*. However, many real-world manifolds do not admit a single global group action, particularly when the topology is complex or the symmetry transformations are only valid patchwise.

To accommodate these more general situations, one turns to *Lie pseudogroups*, which describe symmetries as *local* invertible transformations defined on overlapping patches rather than a single global group. While each local patch may still look like a quotient of a (finite- or infinite-dimensional) group by a subgroup, these patches need not glue together to form a single, globally transitive space. Instead, one obtains a “locally homogeneous” structure that can be formalized through G-structures, Cartan geometry, or (*G, X*)-structures. In these frameworks, each region of the manifold locally resembles a homogeneous model G/H, with transition functions lying in G. The result is that locally, one retains much of the classical Lie-group geometry. In contrast, globally, the manifold’s topology can exhibit a richer or more complicated form than a single homogeneous quotient.

Finally, while certain finite-dimensional Lie pseudogroups are well-understood—particularly those arising from classical, well-studied Lie groups and geometries—there is no complete classification of all finite-dimensional Lie pseudogroups in full generality. Furthermore, restricting to a finite-dimensional Lie pseudogroup may be inadequate to “walk” through every possible configuration space of a given dimension, particularly when the space is topologically or geometrically complex. In such cases, one typically needs an infinite-dimensional pseudogroup (or its associated groupoid of *germs*) to achieve the full local-to-global flexibility required for navigating these manifolds.

#### A.2.3 Important Theorems for pseudogroup Actions

##### Finite-Dimensional Lie Groups

###### Theorem A.2

(Global Transitivity Implies Homogeneous Space). Let *G* be a finite-dimensional Lie group acting *transitively* on a connected manifold *L*. Then *L* can be identified with the homogeneous space *G/H*, where *H* is the stabilizer (isotropy subgroup) of a chosen point in *L*. Consequently,

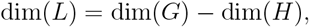

and *L* inherits the topology (and a compatible smooth structure) of the coset space *G/H. Remark*. This classical result underpins much of Lie group geometry and the Erlangen Program. However, many real-world manifolds fail to admit a single globally transitive group—particularly when the underlying space is topologically nontrivial or the relevant symmetry transformations apply *only locally*.

##### Local Neighborhood Results

*Proposition* 2 (Finite Products Span a Connected Lie Group^1^). Let *G* be a *connected* Lie group and *U* ⊆ *G* be a neighborhood of the identity. Define

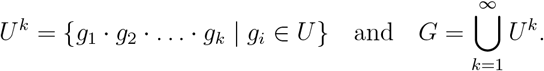

Then every element *g* ∈ *G* can be written as a finite product of elements from *U*. In particular, a sufficiently small neighborhood of the identity can generate the entire group *G* via multiplication.

*Remark*. For *connected, compact* matrix Lie groups, the exponential map is even surjective,^75^ ensuring every group element arises from exp(g). This link between local (near the identity) and global structure also appears in the equivalence of a local Lie group and its Lie algebra [1, Theorem 1.54]. This theorem shows that smoothness implies that the behavior of the group is already specified in any small patch containing the identity. In a sense, Lie groups are “cyclical”: all the elements can be obtained by repeated action of a subset.

##### Local Transitivity on Manifolds

###### Proposition 3

(Local Transitivity in a Neighborhood). Let *C* be a smooth manifold of dimension *n*. For each point *p* ∈ *C*, there exists a neighborhood *U*_*p*_ ⊂ *C* and a local Lie group *G*_*p*_ ⊆ *GL*(*n*, ℝ) acting *transitively* on *U*_*p*_.

*Sketch*. Around each *p*, the tangent space *T*_*p*_*C* is an *n*-dimensional vector space. A sufficiently small subgroup of *GL*(*n*, ℝ) acts transitively on this tangent space by matrix multiplication, sending any vector to any other. By the usual “exponential flow” or “local flow-box” arguments, one obtains a local transitive action on a neighborhood *U*_*p*_ ⊂ *C*.

##### From Lie Groups to Lie Pseudogroups

In many real-world applications, *no single global* finite-dimensional group acts transitively on the entire manifold. One thus moves to *Lie pseudogroups*, which allow *local*, overlapping symmetries that need not glue into a single global group.

###### Theorem A.3

(Transitivity of a Lie Pseudogroup^161^). Let 𝒢 be a Lie pseudogroup acting on a connected manifold *M*. Suppose 𝒢 is *locally transitive* (i.e., for every *p* ∈ *M*, there exists a neighborhood *U*_*p*_ ⊂ *M* where 𝒢 acts transitively). Then 𝒢 is *globally transitive* on *M*.

*Sketch*. The connectedness of *M* and the closure properties of 𝒢 (under composition and inversion) ensure that local transitivity patches extend across overlapping neighborhoods. Iterating these transformations chains together local symmetries to cover the entire manifold, yielding global transitivity.

##### Homogeneous Spaces vs. Local Homogeneity

When a *global* Lie group *G* acts transitively on *M*, we get the classical homogeneous-space identification *M* ≅ *G/H*. By contrast, in the pseudogroup setting, each neighborhood may locally resemble a homogeneous model 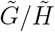, but these local models need not glue consistently into a single global quotient. Instead, one obtains a *locally homogeneous* structure formalizable via *G*-structures or Cartan geometry.^162,163^

##### Dimension and Classification

- **Finite-Dimensional Pseudogroups:** Certain classes of finite-dimensional Lie pseudogroups (e.g., isometry groups, projective groups) are well-studied. However, there is *no complete classification* of all finite-dimensional Lie pseudogroups, reflecting their richness and the vast variety of PDEs that can define them.
- **Infinite-Dimensional Pseudogroups:** In highly complex or topologically intricate manifolds, one typically needs *infinite-dimensional* pseudogroups (or their groupoid of germs) to navigate the space fully.

In short, finite-dimensional groups or pseudogroups may suffice on relatively simple or homogeneous manifolds. But for general manifolds—especially those arising in real-world applications—*locally* defined infinite-dimensional pseudogroups often provide the necessary flexibility to account for patchwise symmetries, reflecting the manifold’s richer topology and geometry.

#### A.2.4 Action of a Lie group on a manifold

The implications of Lie groups being generated by “repeating an operation many times” are profound. They imply that group elements can be written as exponentials of operators. Such exponential operators, in turn, have important properties.

For example, if *f* (*z*) is invariant under an infinitesimal transformation (1 + *ϵT*), so that

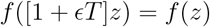

then for small *ϵ*, the function *f* does not change to first order in *ϵ*. This implies that the Lie derivative of *f* with respect to the generator *T* is zero, ℒ_*T*_ *f* = *T* · ∇*f* = 0. If this invariance holds for all infinitesimal transformations, then by repeating the process infinitely many times, which can be conceptualized as taking the limit as *ϵ* becomes infinitesimally small and the number of applications goes to infinity, one would see *f* to be invariant under the finite transformation generated by exponentiating *T*, that is *e*^*ϵT*^ :

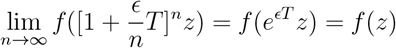

More generally,

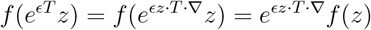

The second equality can be checked using a Taylor expansion or can be justified by checking it to first order and using the limit expression of the exponential as a repeated action of the operator. The first, which provides the connection with the tangent space of the Lie group, follows trivially from 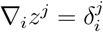, so that *ϵz* · *T* · ∇*z* = *ϵz* · *T*.

Thus, one can inspect the behavior of a function under infinitesimal transformations to infer some properties of the finite ones. Let 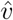 be the operator associated with the vector field *v*, as in *v* = Σ ^*n*^ *ξ*^*n*^∂*/*∂*z*_*n*_, then the Lie series of a function *f* (*z*) at a point *z* can be expressed as:

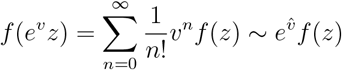

where 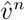 denotes the *n*-th power of the Lie derivative operator 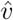 acting on the function *f*, and the symbol ∼ denotes that the series on the right is the Lie series expansion of the left-hand side with some abuse of notation.

The 1D case is a special instance where the manifold is ℝ, and the Lie group is the group of translations on ℝ. The Lie algebra element is 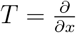, and the exponential map corresponds to adding *λ* to the argument of *f* :

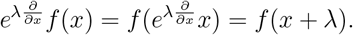

This is the standard Taylor series expansion of *f* about *x*, which is the flow of *f* under the action of the translation group.

#### A.2.5 Invariance and equivariance

To describe the coordinated behavior of transformations induced by a group on different spaces, the terms *invariance* and *equivariance* are often used. The latter is the most general one, indicating that objects in different spaces transform appropriately under the action of the group. Specifically, they may carry different representations, but a commutative diagram linking the actions of the group in both scenarios exists. This can be represented as a commutative diagram,

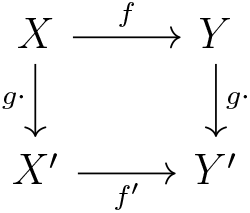

In the diagram, *X* and *Y* are spaces, *f* is a function from *X* to *Y*, and *g* represents the group action. The transformed spaces and functions are denoted by *X*^′^ and *f* ^′^ respectively. The diagram commutes, meaning that starting from any object and following any path through the diagram results in the same outcome.

In more detail, for a function *f* : *X* → *Y* and a group action *g*, the function *f* is said to be *equivariant* under the action of the group if the following relationship holds:

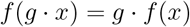

This means that applying the group action *g* to the input *x* and then applying the function *f* is equivalent to first applying the function *f* to *x* and then applying the group action *g* to the result.

For a function *f* : *X* → *Y* and a group action *g*, the function *f* is said to be *invariant* under the action of the group if the following relationship holds:

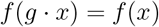

This means that applying the group action *g* to the input *x* does not change the output of the function *f*. In other words, the function’s value remains unchanged or “invariant” under the transformation. It is worth noting that invariance can be considered a special case of equivariance where the transformation on the output space is the identity operation.

#### A.2.6 Further Notes on Lie Groups

The original inspiration sources for Lie were differential equations. But this is now unfortunately mostly forgotten:^1^ the associated Lie groups are not particularly elegant. E.g., a) non-semi-simple nor solvable, b) typically acting nonlinearly on the underlying space, outside (linear) representation theory, and c) may even be only locally defined near identity.

The full range of applicability of Lie groups to differential equations is yet to be determined.^1^

Lie discovered that for continuous groups, the complicated nonlinear conditions of invariance of a system under group transformations can be replaced by equivalent but far simpler linear conditions associated with infinitesimal transformations under the generators of the group.^1^

Infinitesimal symmetry conditions—the defining equations of the symmetry group—can usually be solved.^1^ Once this is done, one can a) construct new solutions from known ones (this provides the means for defining equivalent classes of solutions), b) classify families of differential equations that depend on parameters or other form factors.^1^

Normally, all other things being equal, one seeks the most symmetric equations possible (simplicity).^1^

In the case of ODEs, invariance under a one-parameter symmetry group implies that the order of the ODE can be reduced by one (in the first-order case, it can be solved by quadrature). Multiparameter symmetry engenders further reductions in order, but quadrature alone is not sufficient to solve them.^1^

Symmetry can lead to unique bifurcation phenomena. Hopf bifurcations in symmetric systems can result in synchronized oscillations among a group of neurons due to symmetry constraints. This synchronization can be crucial for understanding phenomena like coherent neural oscillations in the brain.

In essence, the presence of symmetry in neural networks can impact the conditions and characteristics of Hopf bifurcations. The idea is that the network’s symmetries play a role in shaping the bifurcation phenomena and the resulting synchronized oscillations observed in neural systems. The exact mathematical details of how symmetries influence Hopf bifurcations would depend on the specific network structure and equations governing the neural dynamics.

##### Degeneracy of Eigenvalues and Symmetry

When a dynamical system possesses certain symmetries, it can constrain the eigenvectors of the Jacobian matrix A. Symmetry can force some of the eigenvalues of A to be degenerate, meaning they have repeated values. The reason for this degeneracy lies in the symmetry-induced constraints on the dynamics. Symmetry often implies that there are multiple equivalent directions or modes of perturbation around an equilibrium point. These equivalent directions correspond to eigenvectors associated with the degenerate eigenvalues. Mathematically, the degenerate eigenvalues represent directions in which the linearized system’s behavior is indeterminate due to the symmetry-related constraints. They indicate that there are multiple possible linear combinations of these eigenvectors that result in equivalent perturbations. Eigenvalue degeneracy can be associated with bifurcation points where the stability of the equilibrium point changes, and new dynamical behaviors emerge as system parameters vary. The degeneracy of eigenvalues can lead to complex dynamics, such as limit cycles or pattern formation.

It is important to distinguish symmetry in the equations from symmetries in the state or the solution, which may be affected by asymmetry in the initial conditions or external inputs (*symmetry breaking*).

##### Symmetry in Equations (Mathematical Symmetry)

This refers to the inherent symmetries or invariances present in the mathematical equations that describe the system’s dynamics. These symmetries are independent of initial conditions (ICs) or external inputs and are related to the structure of the equations themselves. For example, if the equations exhibit translational symmetry, it means that the system’s behavior remains the same when spatial coordinates are shifted.

##### Symmetries in the State or Solutions (Physical Symmetry)

This relates to the actual configurations or states that the system can adopt as solutions to the equations. These symmetries are observable and depend on the initial conditions, external inputs, and the system’s dynamics. Symmetry in the state implies that the observable patterns or configurations of the system exhibit certain symmetrical properties. These symmetries can be influenced by asymmetries in ICs or external inputs.

The key point is that while the equations themselves may possess symmetries, the actual behavior or state of the system may or may not exhibit these symmetries, depending on the specific conditions and inputs. External factors, such as uneven initial conditions or spatially varying external inputs, can break the symmetry of the system’s state, leading to the emergence of patterns and symmetry-breaking phenomena like Turing bifurcations.

**Symmetry Breaking** is a fundamental concept in understanding how complexity and spatial patterns emerge in natural systems, particularly in the context of spatially extended systems and phenomena like Turing bifurcations. It refers to a phenomenon where the physical state or solutions of a system deviate from or do not exhibit the symmetries that are present in the mathematical equations governing the system. In classical systems^b^, symmetry breaking can arise due to the interplay between asymmetric initial conditions (ICs) or external inputs and the system’s response to these factors. This response is often governed by intrinsic nonlinear dynamics within the system. However, it is crucial to note that the nonlinearities in the equations themselves do not inherently break symmetry; rather, they can amplify or manifest asymmetries introduced by the initial conditions or external inputs.

#### A.2.7 Finite-Dimensional Lie Groups vs. Lie Pseudogroups and Moduli Stacks

In the main text, we emphasize that generative models are best described using *Lie pseudogroups*, which handle local symmetries on overlapping patches of a manifold.^161,164^ However, to appreciate the full generality and power of this approach, it is beneficial to relate it to the classical framework of finite-dimensional Lie groups and the modern concept of moduli stacks.^165^

##### Classical Lie Groups and Homogeneous Spaces

When a finite-dimensional Lie group *G* acts *transitively* on a connected manifold *L, L* can be identified with the homogeneous space *G/H*, where *H* is the stabilizer subgroup of a chosen reference point. This identification constrains the dimension and topology of *L*, with dim(*L*) = dim(*G*) − dim(*H*), and *L* inherits the smooth structure of the coset space *G/H*.^1,75^ However, many real-world configuration spaces do not admit such globally transitive group actions, especially when dealing with complex topologies or localized symmetries.

##### Lie Pseudogroups and Moduli Stacks

To address these complexities, we adopt the framework of Lie pseudogroups, which allow for locally defined symmetry transformations that do not extend to a single global group action. This local flexibility is essential for modeling configuration spaces with intricate or non-uniform symmetry structures. The appropriate mathematical language to encapsulate this setup is that of *moduli stacks*. A moduli stack ℳ = [*X/𝒢*] represents the quotient of a configuration space *X* by a Lie pseudogroup 𝒢, retaining detailed information about local stabilizer subgroups and ensuring that the manifold’s topology and geometry are accurately captured.

##### Implications for Neural Networks and Symmetry Learning

By framing configuration spaces as moduli stacks, we recognize that symmetries in data are often local and may involve complex stabilizer structures. Consequently, neural networks or agents tasked with learning such data must internalize these local symmetries to generalize effectively. This necessitates that network parameters or architectures implicitly encode the transformations corresponding to these symmetries, whether through weight-sharing schemes, equivariant layers, or other structural constraints. The moduli stack framework thus provides a rigorous foundation for understanding how and why neural networks must learn and represent a wide range of symmetries to perform robustly across diverse data transformations.

##### Key Theorems

###### Theorem A.4

(Local Transitivity and Moduli Stacks). Let 𝒢 be a Lie pseudogroup acting on a connected manifold *M*. If 𝒢 is locally transitive (i.e., for every point *p* ∈ *M*, there exists a neighborhood *U*_*p*_ ⊂ *M* where 𝒢 acts transitively), then the configuration space *M* can be described as a moduli stack [*M/𝒢*]. Each local patch *U*_*p*_ is modeled by a homogeneous space *G*_*p*_*/H*_*p*_, where *G*_*p*_ is a local Lie group and *H*_*p*_ is its stabilizer subgroup.

*Sketch*. Given the local transitivity of 𝒢, each neighborhood *U*_*p*_ around a point *p* ∈ *M* resembles a homogeneous space *G*_*p*_*/H*_*p*_. The moduli stack [*M/𝒢*] effectively stitches together these local homogeneous models, preserving information about the local stabilizers and ensuring that the global topology of *M* is accurately represented.

*Proposition* 4 (Finite Products Span a Connected Lie Group^1^). Let *G* be a connected Lie group and *U* ⊆ *G* a neighborhood of the identity. Define

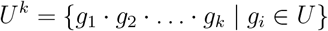

for each *k* ∈ ℕ. Then

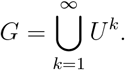

*Remark*. This proposition illustrates that finite-dimensional Lie groups can be generated locally by small neighborhoods around the identity. However, extending these local transformations to a global action on a complex manifold often requires the richer structure provided by Lie pseudogroups and moduli stacks.

*Proposition* 5 (Local Transitivity in a Neighborhood). Let *C* be a smooth manifold of dimension *n*. For each point *p* ∈ *C*, there exists a neighborhood *U*_*p*_ ⊂ *C* and a local Lie group *G*_*p*_ ⊆ *GL*(*n, ℝ*) acting transitively on *U*_*p*_.

*Sketch*. In the vicinity of any point *p*, the tangent space *T*_*p*_*C* is *n*-dimensional, allowing the local Lie group *G*_*p*_ to act transitively via linear transformations. Using the exponential map and flow-box arguments, these local actions extend to a neighborhood *U*_*p*_, ensuring transitivity within *U*_*p*_.

###### Theorem A.5

(Transitivity of a Lie Pseudogroup^161^). Let 𝒢 be a Lie pseudogroup acting on a connected smooth manifold *M*. If 𝒢 is locally transitive (i.e., for every *p* ∈ *M*, there exists a neighborhood *U*_*p*_ ⊂ *M* where 𝒢 acts transitively), then 𝒢 is globally transitive on *M*.

*Sketch*. The connectedness of *M* and the closure properties of 𝒢 ensure that local transitivity can be extended across overlapping neighborhoods. By chaining local transformations, 𝒢 achieves global transitivity.

###### Conclusion

While finite-dimensional Lie groups provide a powerful framework for understanding symmetries in homogeneous spaces, their global applicability is limited in complex or topologically intricate manifolds. Lie pseudogroups, complemented by the moduli stack formalism, offer a more flexible and comprehensive approach to modeling and analyzing symmetries in such settings. This enriched language not only captures local symmetries with finite-dimensional parameters but also integrates seamlessly with the nuanced topology of real-world configuration spaces, making it an indispensable tool for studying neural network symmetries and their structural implications.

### A.3 Groups, Turing Machines, and Generative Models

In the discrete context, the analog of Lie groups is finitely generated groups, where a finite set of generators produces all elements, with cyclic groups representing cases generated by a single element. Here, we discuss the idea that Turing machines implicitly use groups, or some relaxed versions of them such as *monoids*,^166^ for computation.

Can we think of Turing machines, and hence computation, as a branch of group theory? While the practicality and exact mapping pose significant challenges, the conceptual framework offers intriguing possibilities for future research in computational theory and artificial intelligence.

Group theory, a branch of abstract algebra, deals with the study of groups that are sets equipped with a binary operation that satisfies certain axioms: closure, associativity, identity, and invertibility. This mathematical framework can be applied to studying computational models, such as Turing machines, particularly in understanding the symmetry and structure of computational processes.

To begin analyzing this, we will use the three-tape Turing machine^167^ with a finite private tape, which can be conceptualized as a generative model. In this model, the machine’s states and the states of the private tape could be interpreted as elements of a group. The transition from one state to another, governed by the machine’s rules and the symbols on the tapes, can be analogized to a group operation.

A three-tape Turing machine is an extension of the standard Turing machine model with three separate tapes: the input tape, the output tape, and a private (or work) tape. Each tape has its own tape head for reading and writing. It can be formally defined as a 7-tuple (*Q*, Σ, Γ, *δ, q*_0_, *q*_accept_, *q*_reject_) where:

- *Q* is a finite set of states.
- Σ is a finite input alphabet that does not contain the blank symbol ⊔.
- Γ is the tape alphabet, where ⊔ ∈ Γ and Σ ⊆ Γ.
- *δ* : *Q* × Γ^3^ → *Q* × Γ^3^ × {*L, R, S*}^3^ is the transition function. Here, *L, R*, and *S* denote left shift, right shift, and no shift on the tapes, respectively.
- *q*_0_ ∈ *Q* is the initial state.
- *q*_accept_ ∈ *Q* is the accept state.
- *q*_reject_ ∈ *Q* is the reject state, distinct from the accept state.

The machine operates as follows:

- The input tape contains the input string and is read-only.
- The output tape is used to write the output and is write-only.
- The private tape is used for intermediate computations and can be both read from and written to.
- The transition function *δ* dictates the machine’s actions based on the current state and the symbols under the tape heads. It specifies the next state, the symbols to write on each tape, and the movements of the tape heads.
- The computation begins in the initial state *q*_0_ and proceeds according to the transition function until the machine enters either the accept state *q*_accept_ or the reject state *q*_reject_.

The challenges in mapping Turing machines to group theory are

- **Closure:** Ensuring that the combination of any two states (or actions) results in another valid state within the system.
- **Associativity:** The sequence of transitions must not affect the final state of the machine, a non-trivial property to verify in computational processes.
- **Identity and Invertibility:** Identifying a state that acts as an identity element, and for each state, an inverse state, can be complex in the context of a Turing machine.

The minimal requirement is that of composition. We can relax the other two, and instead of groups, we have more general algebraic structures. A *magma*, or *groupoid*, is a foundational concept where a set is equipped with a binary operation, with the only requirement being closure. An extension of this is a *semigroup*, which is an associative magma, meaning the binary operation is associative. Building upon this, a *monoid* is essentially a semigroup that includes an identity element but does not necessarily have inverses for all its elements. Finally, a *group* is a monoid where every element has an inverse, thus completing the hierarchy from the most general structure (magma) to the more specialized one (group). While the concepts of Lie algebras and exponential maps are not directly applicable to magmas, semigroups, and monoids in the way they are to Lie groups, there are efforts and research in mathematics to explore analogous or related structures in these broader contexts.

For example, the concept of monoids becomes necessary when dealing with computational systems where inverses are not available, such as cellular automata like Rule 110.^168^ A monoid is an algebraic structure similar to a group but does not require every element to have an inverse. In Rule 110, the evolution operator *T* that updates the cellular configuration lacks an inverse because the process is generally irreversible—the previous state cannot be uniquely determined from the current state.

The power of recognizing the monoid structure in Rule 110 lies in understanding how the repeated application of a simple rule can generate highly complex behavior. Despite the simplicity of the rule (a single generator *T*), the iterative process *T*^*n*^ produces rich dynamics that are capable of universal computation (Rule 110 is Turing complete). This highlights that even without the full framework of group theory, the presence of an underlying algebraic structure like a monoid is sufficient to capture the essence of the system’s generative process.

### A.4 Symmetry in ODEs: abstract vs. traditional definitions

Symmetry plays a fundamental role in the analysis and solution of ordinary differential equations (ODEs). In this section, we present an abstract definition of symmetry based on transformations of initial conditions (ICs) and compare it with the traditional definition that involves transformations of variables. We demonstrate that both definitions are equivalent in their ability to map solutions to solutions while preserving the form of the ODE and explore the structure of the resulting symmetry groups, highlighting that they can be either abelian or non-abelian, depending on the system.

#### A.4.1 Olver’s definition of symmetry of PDEs^1, 2^

Consider a general system of *n*^th^ order (partial) differential equations

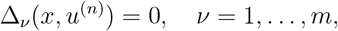

in *p* independent variables *x* = (*x*^1^, …, *x*^*p*^), and *q* dependent variables *u* = (*u*^1^, …, *u*^*q*^), with *u*^(*n*)^ denoting the derivatives of the *u*’s with respect to the *x*’s up to order *n*. In general, by a *symmetry of the system (1)*, we mean a transformation that takes solutions to solutions. The most basic type of symmetry is a (locally defined) invertible map on the space of independent and dependent variables:

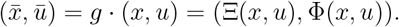

Such transformations act on solutions *u* = *f* (*x*) by pointwise transforming their graphs; in other words, if Γ_*f*_ = {(*x, f* (*x*))} denotes the graph of *f*, then the transformed function 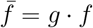 will have the graph

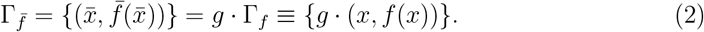

##### Definition A.6.

A local Lie group of transformations *G* is called a *symmetry group* of the system of partial differential equations (1) if 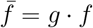 is a solution whenever *f* is.

##### A.4.2 Abstract Definition of Symmetry for ODEs

Here, we provide a generalized definition of symmetry. As it turns out, it is closely aligned with that of Olver for PDEs.^1,2^ Consider an autonomous ODE of the form:

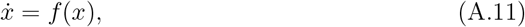

where *x* ∈ ℝ^*n*^ is a vector of dependent variables, and *f* : ℝ^*n*^ → ℝ^*n*^ is a smooth function.

Let *x*(*t*; *x*_0_) denote the unique solution of the ODE with the initial condition *x*(0) = *x*_0_. The mapping from the initial condition *x*_0_ to the solution *x*(*t*; *x*_0_) establishes a one-to-one correspondence between initial conditions and solutions due to the existence and uniqueness theorems for ODEs.

**Definition (abstract definition of symmetry):** A *symmetry* of the ODE is any **bijective transformation** *γ* acting on the initial conditions *x*_0_, such that:

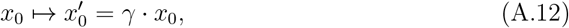

which induces a transformation of the solution:

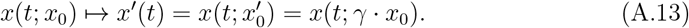

Since each initial condition corresponds to a unique solution, any bijective transformation *γ* on *x*_0_ automatically maps a solution to another solution that satisfies the same ODE,

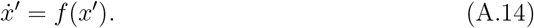

Thus, the transformation *γ* preserves the form of the ODE by ensuring that *x*^′^(*t*) satisfies the original differential equation.

The set of these transformations forms a Lie group, the group of diffeomorphisms of the manifold of initial conditions. Under some conditions, the Lie group has a finite number of generators.

#### A.5 Are all Lie generative models finite-dimensional?

Suppose we have a generative model for some dataset. This means that we can generate variants of the same object in the dataset. The dataset itself or its model can both be seen as that which remains constant, e.g., “catness”. Thus, we talk about *symmetry, continuous transformations* (perhaps discrete ones, too) that leave the essence of the object (*catness*, or the dataset) invariant. We also expect that it is possible to rewrite the generative model using any coordinate system: there is no particular choice of coordinates in configuration space that is better than others, there is no particular origin, and no special cat.

The notion that a generative model can be described by a finite Lie group highlights the importance of symmetry and smoothness in modeling natural data. When a model exhibits these properties, it allows us to understand and navigate the configuration space in a structured and efficient manner, which is particularly useful in practical applications.

If natural data can be captured in a latent space—a lower-dimensional representation where complex relationships are simplified—we can choose an arbitrary point in this latent space as a reference. This is the case with neuroimaging data, which includes complex brain activity patterns obtained from techniques like fMRI or EEG. Embedding such data in a latent space (e.g., processed using a variational autoencoder^112^) reveals underlying structures and patterns not immediately apparent in the high-dimensional raw data. By studying the relationship between displacements in different directions around the reference point in the latent space, we can gain insights into how variations in the data correspond to meaningful changes in brain activity. If this can be defined by a Lie group (at least locally), then this relationship should be independent of the choice of the reference point. For example, the structure of the group rotations in the latent space of cat images should act independently of the choice of the reference cat.

We can refine the requirements for associating such a group with a generative model as follows. Let *C* be a smooth manifold. We say that *C* admits a *point-transitive finite-dimensional Lie subgroup* of Diff(*C*) if there exists a finite-dimensional Lie subgroup *G* ⊆ Diff(*C*) such that for any two points *x, y* ∈ *C*, there exists *g* ∈ *G* such that *g* · *x* = *y*. In the case of *1D manifolds*, every connected, smooth manifold is diffeomorphic either to R or to *S*^1^, the circle. For both these types, there exists a finite-dimensional Lie subgroup of the diffeomorphism group that acts point-transitively. Specifically, translations generate such a group for R, and rotations for *S*^1^. Thus, any generative model using these 1D manifolds is a Lie generative model.

However, not all configuration space manifolds have the property of finite Lie transitivity. Mostow’s Rigidity theorem^169^ provides insights into point-transitive actions on *surfaces (2D manifolds)*.^170^ According to the theorem, a 2D manifold *C* without boundary admits a point-transitive Lie group action if and only if *C* is a plane, sphere, cylinder, torus, projective plane, Möbius strip, or Klein bottle. As we move to higher-dimensional manifolds, a comprehensive classification similar to Mostow’s theorem becomes increasingly complex. Nevertheless, certain partial classifications and criteria exist that specify the conditions under which a manifold allows a transitive Lie group action.

For hyperbolic surfaces of genus *g >* 1, no Lie group exists that can act transitively on the entire manifold. This reflects the lack of global symmetry inherent to such spaces, which contrasts with the high degree of symmetry seen in spherical or flat geometries where Lie groups can act transitively. While these manifolds are locally homogeneous, their global structure breaks this symmetry, making them navigable only through local actions described by pseudogroups. Mostow’s Rigidity Theorem reinforces this insight by showing that the geometry of hyperbolic manifolds is uniquely determined by their fundamental group, which does not admit the symmetry required for a transitive Lie group action.

Despite the difficulties in finding the conditions for the existence of global finite Lie groups acting transitively on a given manifold, transitivity can be expected in local Lie group action. The proof for *global* transitivity requires specific properties or structures on *C*. Establishing global transitivity is equivalent to showing that for any two points *c*_1_, *c*_2_ ∈ *C*, there exists an element in the Lie group that can transform *c*_1_ into *c*_2_. Mostow’s theorem suggests there are deep links between the geometry of C and the algebraic structure of groups acting transitively on it.

Mostow’s theorem suggests wider deep links between the geometry of a manifold 𝒞 and the algebraic structure of groups acting transitively on it. When a Lie group *G* acts transitively on a manifold 𝒞, the manifold can often be described as a *homogeneous space* 𝒞 ≅ *G/H*, where *H* is the stabilizer subgroup of a point in 𝒞. The topology of 𝒞 is intimately connected to the topology of *G* and *H*. For example, the dimensionality of 𝒞 corresponds to the co-dimension of *H* in *G*, while the connectedness of 𝒞 reflects the transitivity of the connected component of the identity in *G*. Moreover, the geometry of 𝒞, such as its curvature, imposes constraints on the structure of *G*, influencing properties like compactness and the existence of certain representations.^170^

For pseudogroups, which generalize the concept of Lie group actions to local transformations, the relationship becomes more nuanced. A pseudogroup can act locally on C, adapting to its geometry in ways that global Lie group actions cannot. The topology of C interacts with the infinitesimal generators of the pseudogroup, which are often solutions to partial differential equations governing local symmetries.^1,161^ Unlike global actions, pseudogroups are particularly suited to manifolds with local symmetries, such as foliated manifolds or those with varying curvature.

These relationships illustrate the profound interplay between algebra and geometry, where the structure of 𝒞 constrains the algebraic properties of *G*, and for pseudogroups, influences the nature of local transformations. This connection underscores the central role of symmetry in understanding both global and local manifold properties, providing a bridge between differential geometry and algebraic structures.

#### A.6 Conserved quantities and canonical variables

In this section, we provide some background on the theory of symmetries and constraints in ODEs. Let us consider, for simplicity, a well-behaved autonomous system of *X* equations and unknowns,

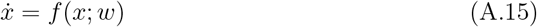

(with a smooth function *f*). The solution involves *X* integration constants. Paraphrasing the classical argument in Landau and Lifschitz,^171^ since the equations of motion for such a closed system do not involve time explicitly, the choice of the origin of time is entirely arbitrary, and one of the arbitrary constants in the solution of the equations can always be taken as an additive constant *t*_0_ in the time, and the solution written as *x* = *x*(*t* + *t*_0_, *C*), with *C* ∈ ℝ^*X*−1^.

We can use one of these *X* equations to (at least locally) express *t* + *t*_0_ as a function of *x* and the rest of the constants. We can think of the corresponding component of *x*, say *x*_0_, as the “clock” (this is actually what real clocks are)^c^, and we write *t* + *t*_0_ = *u*(*x*_0_, *C*). Substituting this, the solution can then be written as *x*_*n*_ = *x*_*n*_(*x*_0_, *C*) for the other *X* − 1 components.

Thus, we have *X* − 1 constraints,

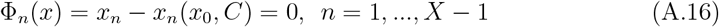

that the solutions must satisfy.

Since solutions cannot cross, one may expect a one-to-one correspondence between *C*’s and solutions. However, this may fail when the constants of the motion are implicitly defined through non-injective functions. In cases where a constant of motion *Q*(*x*) is expressed in terms involving non-injective functions, such as *Q*(*x*) = sin(*C*), the map from solutions to constants of motion is no longer one-to-one, leading to multivalued mappings.

Up to this multivalued ambiguity, we can now use the *X* − 1 equations in A.16 to express the remaining *C*s as a function of *x*. We thus obtain *X* − 1 independent constants of the motion,

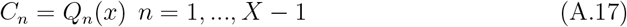

so 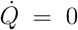. We may rewrite this more rigorously by choosing a fixed branch of the solution.

Therefore, any given solution in *X* dimensional space is constrained by *X* − 1 conserved quantities^d^, which results, as we already knew from the start, in one-dimensional trajectories.

There cannot be any further independent conserved quantities unless the solution is a fixed point, but any combination of constants of the motion is a constant of the motion.

While the autonomous system of equations defined by 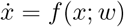, where *f* is a smooth function, allows us to find *X* − 1 conserved quantities *Q*_*n*_(*x*), it is crucial to consider the differentiability of these quantities across the entire phase space. Evaluating the time derivatives of these constants of the motion, we have

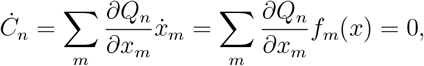

implying the conservation over time. However, this expression entails potential singular points where components of *f* (*x*) vanish or where the partial derivatives 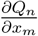 become singular, rendering *Q*_*n*_(*x*) not well-defined. These singular points introduce barriers to implementing a global change of coordinates in phase space using these conserved quantities, as they may not be differentiable at these points.

When it is possible to find such canonical variables globally, the system is called integrable. An *integrable system* is one for which it is possible to find as many independent constants of motion as degrees of freedom, which allows for the system to be solved exactly through analytic methods. These constants of motion are typically associated with symmetries of the system. Integrability is a strong condition and is typically hard to achieve or prove for most systems, especially when going beyond two degrees of freedom.

##### A.6.1 Some examples

Consider the forced non-relativistic particle, 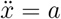, or, in first-order form

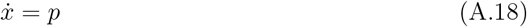

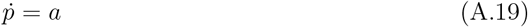

If *x*(*t*) is a solution, then 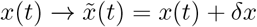 is also a solution. The transformation 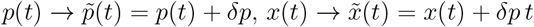, is also a symmetry, since

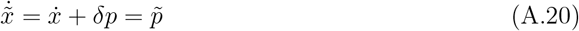

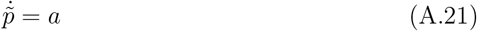

so, the overall symmetry of the ODE system is

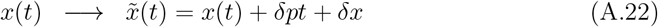

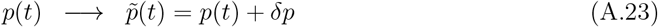

The solution to the equations with *a* = 0 is *p* = *p*_0_ and *x* = *t* + *t*_0_, for arbitrary constants *p*_0_ and *t*_0_. Following the argument above, we have *t* + *t*_0_ = *x* (the *x* coordinate can be used as a natural clock) and the conservation law *C*(*x, p*) = *p* = *p*_0_. The symmetry transformation shifts *C* by *δp*.

It is instructive to consider the case *a≠* 0. Here, we have translational invariance without momentum conservation. Again, we use a variable as a clock, this time *p/a* = *t* + *t*_0_, and express the solution as a function of the clock and constants of integration,

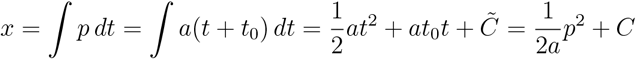

The conserved quantity is 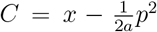. The symmetry transformation shifts *C* by 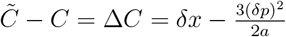.

#### A.6.2 A not-so-simple example of constrained dynamics

Consider the constrained ODE (differential algebraic equation)

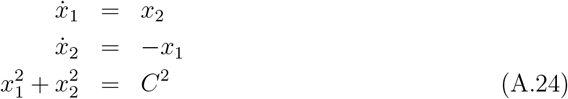

This problem has a solution because the general solution to the first two equations is

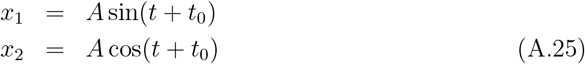

for any constant *A*, which automatically guarantees that 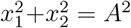. Hence, because the constraint is compatible with the conserved quantities of the ODEs, finding a solution to the three equations is possible. Adding an incompatible constraint, e.g., *x*_1_ = 1, would force fixing the time variable *t* because the intersection of the trajectory with the constraint would be a single point. For more clarity, we may express the system of equations in symmetry manifesting canonical coordinates (polar coordinates, in this case, corresponding to *x*_1_ and *x*_2_),

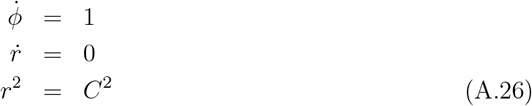

A solution is possible because of the second equation, which manifests the invariance of the ODE component to rescaling of the variables (changes in radius, with generator ∂*/*∂*r*)—translations of *r, r* → *r* + *ϵ*—with its associated conserved variable *r*. Again, the conserved variable corresponds to a center manifold (zero real part eigenvalue of the system).

##### A.6.3 Approximate symmetry after transients

If the second equation were instead 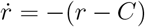, we would have the constraint holding in steady state after transient dynamics,

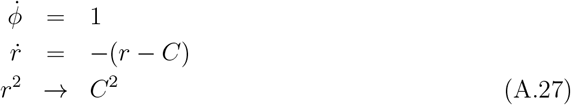

The equations encode the desired constraint value, and they are no longer translational invariant (in *r*). Rather, a simultaneous translation of *r* and *C* leaves the ODE system invariant. We can think of *C* as an input and the equation *r* → *C* as a post-transient constraint. The solution is *r* = *C* + *ae*^−*t*^ → *C*. If *C* changes slowly compared to the transient time scale, the equations will still hold. The input, instead of the initial conditions, implements the group control on the solution space.

This equation has a Lyapunov function, which is also invariant under a simultaneous translation of *r* and *C*, namely 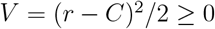, since 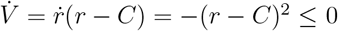.

Another example of how to achieve long-term symmetry (after transients die out) is

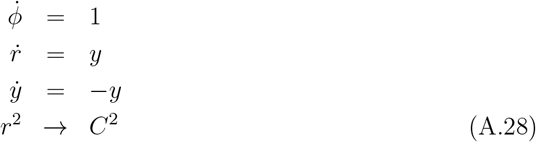

which is equivalent to

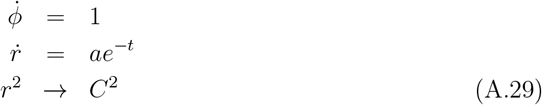

with solution *r* = *C* − *ae*^−*t*^. The trick is to make the equations for *r* evolve in time to the desired form, 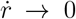, which then delivers the desired properties of symmetry and conservation law. In this case, unlike the previous one, the constraint satisfied is a function of the initial conditions. The Lyupanov function is *V* = *r*^2^*/*2 ≥ 0, since 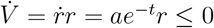 for *a <* 0.

##### A.6.4 Transition to Canonical Coordinates

To advance our understanding of the relationship between symmetry and conserved quantities, we ignore the difficulty of differentiability and assume we can transition to a local “canonical” coordinate system, where each coordinate but one (*x*_0_(*t*)) remains constant, corresponding to the conserved quantities identified previously. Representing these coordinates as *Q*_*i*_, with *i* ranging from 1 to *X* −1, we have the transformed system represented as

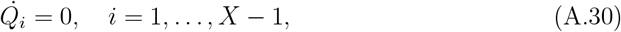

with only one active coordinate evolving with time. The solutions are *x*_0_ = *x*_0_(*t*) for some function *x*_0_ and *Q*_*i*_ = *C*_*i*_. Clearly, any transformation that maps a solution to 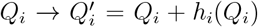 generates a new solution, i.e., is a symmetry of the ODE. The new solutions is simply (*x*_0_(*t* + *t*_0_, *C*^′^), *Q*^′^).

In the canonical coordinates, each conserved quantity engenders a one-parameter symmetry characterized by transformations of the form

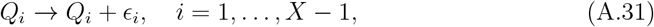

yielding a Lie group with *X* − 1 parameters.

The generators of these symmetries, fundamental to our discussion, can simply be denoted by partial derivative operators with respect to the respective coordinates,

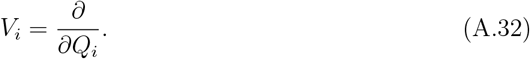

These generators capture the symmetry properties of the system, satisfying commutation relations indicative of the independence of the conserved quantities given by

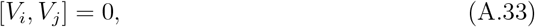

The action of the group on the solution (*x*_0_, *Q*_*i*_) can be described through the transformations (*x*_0_ is left unaffected),

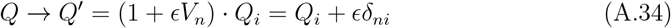

with *δ*_*ni*_ being the Kronecker delta. That is, the effect of the transformation is a shift of the constants of the motion. Note that (*x*_0_, *Q*^′^) is also a solution to the ODE since 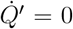. This demonstrates that *V*_*n*_ is the generator of a one-parameter group of symmetries representing translations in the canonical coordinates.

In summary, we established that in a non-trivial *X*-dimensional first-order autonomous ODE, one can exploit time invariance to identify precisely *X* − 1 independent constraints, producing one-dimensional solutions (trajectories). Transitioning to canonical coordinates, when possible, where all but one coordinate (*x*_0_(*t*)) remain fixed, at least locally, we elucidated that symmetry can be engendered through transformations mapping *Q*_*i*_ to *Q*_*i*_ + *h*_*i*_(*Q*_*i*_), yielding new solutions and therefore being symmetries of the ODE system. The generators *V*_*i*_ = ∂*/*∂*Q*_*i*_ foster a group of symmetries encapsulating translations in the canonical coordinates, a Lie group with *X* − 1 parameters.

Finally, our argument shows that there are constraints that lead to “local” conserved quantities and symmetries (and 1D trajectories), but there may not be such globally well-defined quantities. There may be singularities in the conservation functions.^173^

Next, we impose a first hierarchical constraint 𝒞_1_, 𝒞_1_(**y**_1_) = *r* − 1 = 0, which fixes *r* = 1. This constraint reduces the state space to the surface of a unit sphere. The manifold at this level is defined as

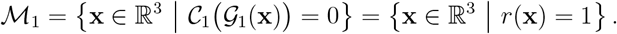

Since we have discarded *ϕ*, the remaining variable is *θ* ∈ [0, *π*]. However, because *ϕ* is not specified, for each value of *θ*, there is a circle of points (a line of latitude) on the sphere. Therefore, our manifold ℳ_1_ corresponds to the set of all such circles on the unit sphere.

## Footnotes

a Real images are digital, and a pixel can take a finite number of values—we ignore such details.

b In quantum systems, quantum fluctuations can also break the symmetry.

c can even add such a coordinate to represent time by adding a corresponding phase space coordinate, e.g., by adding some physical mechanism such as a particle moving in space, so that 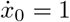 and *x*_0_ = *t* + *t*_0_.

d Integrable systems in the context of differential equations are distinguished by the presence of a full set of *analytic* constants of motion, equal in number to the system’s degrees of freedom172 These constants are not only independent but also in *involution* (i.e., their Poisson brackets vanish), a condition ensuring their functional independence. This concept extends beyond the mere existence of integration constants in solutions to ordinary differential equations, as discussed by Landau. In integrable systems, these analytic constants facilitate a detailed characterization of the system’s dynamics, often allowing the motion to be confined to an invariant torus in the phase space and enabling the system to be solved by quadratures. This precise structure of constants of motion imbues integrable systems with a degree of solvability and predictability uncommon in more general dynamical systems.

